# Pioneering Hybrid Genome Assemblies of *Fusarium graminearum* and *Fusarium avenaceum* from Indian Wheat: Insights into Virulence Frameworks and Genome Evolution

**DOI:** 10.64898/2026.11.18.767454

**Authors:** Govardhan Choppadandi, KJ Yashavanthakumar, Sudhir Navathe

## Abstract

Fusarium head blight (FHB), primarily caused by various species of Fusarium, is a destructive disease of wheat that reduces yield and contaminates grain with mycotoxins. The present study reports high-quality hybrid genome assemblies for two Indian wheat isolates, Fusarium graminearum strain TNW1 and Fusarium avenaceum strain DMW8, representing the first genome-scale comparative study of FHB-associated Fusarium isolates infectious on wheat from India. Both Indian isolates retained conserved Fusarium secondary-metabolite clusters, including aurofusarin, chrysogine, clavaric acid, koraiol, orcinol/orsellinic acid, and oxyjavanicin, while exhibiting distinct species-associated biosynthetic profiles. TNW1 contained zearalenone and DON-related biosynthetic clusters, whereas DMW8 contained beauvericin and other species-associated clusters, highlighting differences in predicted biosynthetic potential between the two Fusarium species. Notably, both genomes showed relatively low transposable-element (TE) content compared with their respective species groups, with TEs occupying 0.96% of the TNW1 genome and 1.55% of the DMW8 genome. RIP-associated genomic coverage was also comparatively low, at 20.01% and 22.38%, respectively. TE insertion analysis identified predominantly reference-associated insertions, with two candidate non-reference insertions detected in DMW8. Gene-family analyses revealed greater duplication and expansion–contraction dynamics in DMW8 than in TNW1. Together, these findings highlight differences in predicted genomic features, secondary metabolite biosynthetic potential, and gene family composition between the two Indian isolates, providing genomic resources for future investigations of Fusarium biology and wheat–pathogen interactions.

## 1. Introduction

Wheat production is affected by various fungal diseases. Among them, Fusarium head blight (FHB), leaf rust (LR), stripe rust (SR), and foliar/blotch diseases such as spot blotch are among the most problematic fungal diseases worldwide, including in India (Singh et al., 2025). Fusarium species cause head blight, also known as scab. The typical symptoms produced by the fungus appear mainly on the glumes and rachis of wheat plants as water-soaked lesions. Later, the fungus spreads within the wheat ear heads, resulting in partial bleaching that progresses to complete blight of the affected ear heads (Wise et al., 2015). First reported in England in 1884, the disease later spread to other parts of the world and now causes damage in humid and semi-humid wheat-growing areas, especially in the USA, Canada, Asia, Europe, Australia, and South America (Dickson et al.,1942). Fusarium head blight significantly reduces grain yield and quality by infecting spikelets or the spike. Infected kernels appear shrivelled and frequently contaminate grain with mycotoxins, which pose serious risks to animals and humans (Selvarajan et al., 2026). The most important toxins produced by *Fusarium* species include trichothecenes, zearalenones, fumonisins, and emerging metabolites such as beauvericin and enniatins (Ferrigo et al., 2016). Contamination of food with mycotoxins has been identified as a global issue in food safety and public health, and toxin production is influenced by climate change and environmental conditions (El-Sayed et al., 2022; Pleadin et al., 2019; Eskola et al., 2020). In wheat–maize cropping systems, *Fusarium* species survive on both wheat straw and maize stubble, driving pathogen adaptation and community shifts (Sun et al., 2025).

In India, the disease was historically considered minor compared to rusts and spot blotch in major wheat-growing zones such as the Northwestern and Northeastern plains zone and parts of the Central Zone (Kaul et al., 2023). However, recent reports indicate its emerging importance in the northern wheat belt and peninsular India, especially under maize rotations and conservation agriculture (Hegde et al., 2023). Despite this, there is limited information on pathogen virulence, distribution, and chemotype shifts. The taxonomy and ecology of the pathogen remain largely unexplored, and molecular data from Indian isolates are scarce in public databases. Owing to diverse climatic conditions and insufficient molecular characterisation, research on this pathogen remains limited in India. Therefore, it is essential to anticipate pathogen adaptation under changing agro-climatic conditions.

Fungal genome studies are very important for understanding the mechanisms of pathogenicity and adaptation in pathogens. Genome analyses enable the identification of virulence-associated genes, including effectors and secreted proteins that manipulate host physiology and suppress plant immune responses, thereby facilitating infection (Lo Presti et al., 2015).

Deoxynivalenol (DON) and the newly identified NX mycotoxins are crucial for *Fusarium graminearum* infection of wheat heads, alongside numerous effector proteins that suppress plant immunity and contribute to Fusarium head blight (Xu et al., 2022; Armer et al., 2024; Rhoades et al., 2024; Li et al., 2022). Similarly, *Fusarium avenaceum* pathogenicity is associated with emerging mycotoxins such as moniliformin, enniatins (A, A1, B, B1), and occasionally beauvericin (Logrieco et al., 2002; Jestoi et al., 2004; Kokkonen et al., 2010; Gautier et al., 2020). In addition, transposable elements (TEs) and TE-rich accessory genomic regions contribute significantly to genome plasticity, driving rapid evolution, host specialisation, and the emergence of new pathogenic variants (López Díaz et al., 2025). Repeat-induced point mutation (RIP), a key fungal genome defence mechanism, further shapes genome evolution by mutating duplicated sequences and influencing the diversification of pathogenicity-related genes (Badet & Croll, 2025).

Before this study, there was no genome-level information available for Indian Fusarium species responsible for Fusarium head blight, and no successful efforts had been reported to sequence their whole genomes or to identify associated virulence genes. To address this gap, we generated high-quality genome assemblies for two Indian *Fusarium* isolates, *Fusarium graminearum* (TNW1) and *F. avenaceum* (DMW8), isolated from infected wheat spikes. We examined genome architecture, transposable elements, repeat-induced mutation, and virulence-related genomic features to better understand how genome structure contributes to evolutionary and pathogenic diversity within the genus.

Considering the growing agricultural impact of FHB, particularly under changing climate conditions and emerging disease pressures, this work focused on isolates infecting spring wheat. Together, these analyses provide foundational genomic resources for Indian isolates and offer new insight into genome plasticity and pathogenic mechanisms in fungal plant pathogens.

## 2. Materials and methods

### 2.1 Sample collection and *Fusarium* isolation

Wheat spike samples showing floret bleaching and tombstone kernel symptoms were collected from Wellington, Tamil Nadu, and Dalang Maidan, Himachal Pradesh, India, in 2024-2025. The fungal species were aseptically isolated on potato dextrose agar (PDA) plates containing 50 μg/mL Tagmycin and incubated at 27±2 °C in the dark for 7–14 days. Suspected *Fusarium-*like colonies were transferred to fresh PDA plates. The monoconidial isolation was performed to obtain pure cultures, which were stored at 27±2 °C for further studies. The pathogenicity of these isolates was tested using susceptible wheat varieties. The isolates were named TNW1 from Wellington and DMW8 from Dalang Maidan.

### 2.2 Morphological characterisation and multi-locus identification of fungal isolates

For morphological identification, the methods described by Summerell et al. (2003) and FusariumID (https://www.fusarium.org/page/Protocols; Crous et al. 2021) were followed. Single-spore isolates were cultured on Potato Dextrose Agar (PDA) and incubated at 25 ± 2 °C for six days. Colony characteristics, including growth rate, pigmentation, and presence of sporodochia, were recorded. For micromorphology, Spezieller Nährstoffarmer Agar (SNA) and Carnation Leaf Agar (CLA) were used.

Genomic DNA was extracted from two isolates at 21 days of mycelial growth, aseptically, in 100ml PD broth at 25 °C, using the CTAB sorbitol protocol described by Inglis et al. (2018) with minor modifications. β-mercaptoethanol and polyvinylpyrrolidone (PVP) were omitted from the sorbitol pre-wash buffer, and mechanical disruption using a bead mill was replaced by manual disruption. Then, DNA concentration and purity were measured using the Thermo Scientific Multiskan SkyHigh spectrophotometer. The integrity of the DNA was observed on an agarose gel (1%).

Molecular identification was performed by PCR amplification of the ITS, TEF-1α, and RPB2 loci (Table 1). PCR products were subjected to Sanger sequencing. The resulting sequences were compared with the curated FUSARIUM-ID database using BLASTn to achieve species-level identification. Further sequences were aligned with additional reference sequences from the NCBI database (see Supplementary File S1) using MAFFT v7.525 and then concatenated using AMAS. Maximum likelihood analysis was conducted in IQ-TREE v2.0.7 (Minh et al. 2020), with the best-fit models selected by ModelFinder based on the Bayesian Information Criterion (BIC). Branch support was estimated with 1000 ultrafast bootstrap replicates.

**Table 1.**
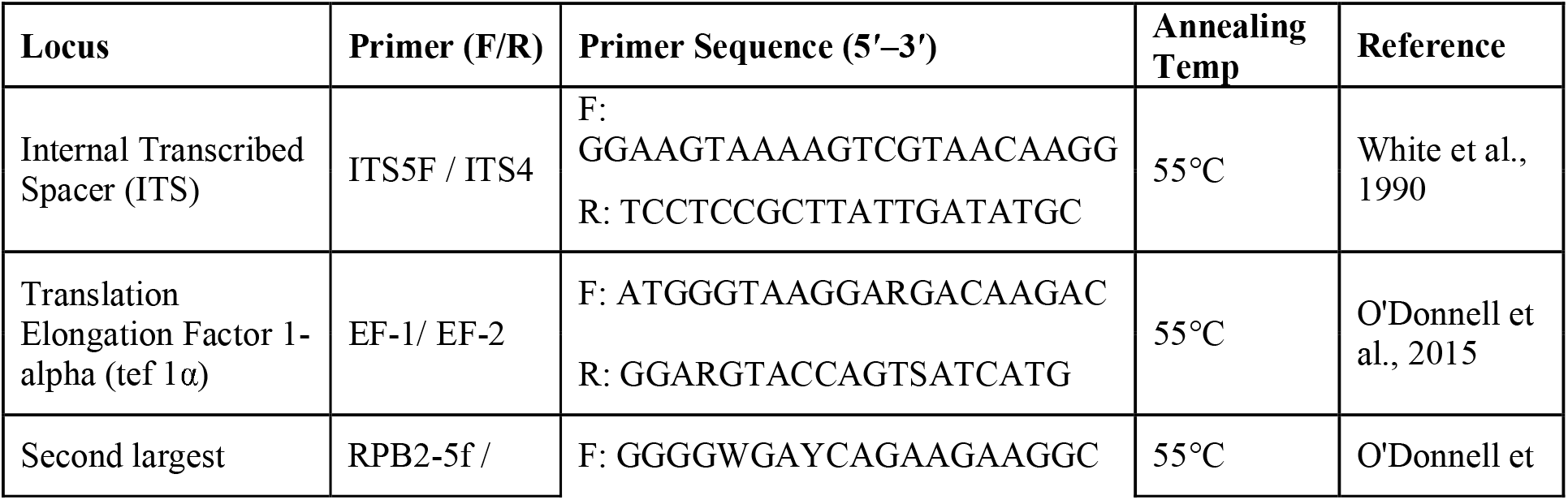
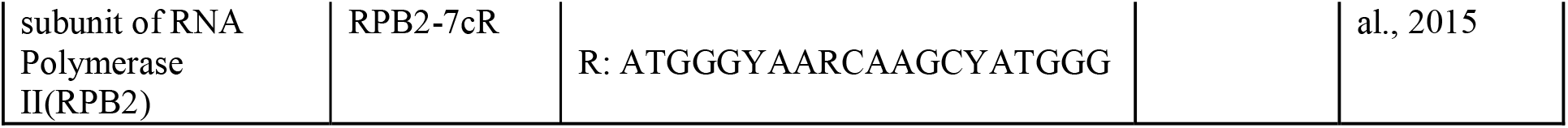
List of primer pairs used for PCR amplification of ITS, *tef1*, and RPB2 gene regions, with their sequences (5′–3′), annealing temperatures, and corresponding references.

### 2.3 Pathogenicity Assay and Disease Assessment

The pathogenicity of *Fusarium graminearum* TNW1 and *Fusarium avenaceum* DMW8 was evaluated on the susceptible wheat genotypes MACS-6222 and MACS-6747 and the resistant genotype Sumai 3 under glasshouse conditions. Non-inoculated MACS-6222 and MACS-6747 plants treated with prothioconazole + tebuconazole to prevent natural FHB infection served as controls. The fungicide was applied at the recommended rate before the anticipated period of infection, and the plants were maintained under fungicide protection until harvest. The fungal isolates were grown in mung bean broth for one week, and the resulting suspensions were filtered through a sterile cotton strainer. Spore concentrations were determined using a Neubauer haemocytometer and adjusted to 1 × 10 spores/mL.

Inoculation was performed either in the early morning, before 06:00 h, or in the evening, after 17:00 h. Cotton wool plugs soaked in the respective spore suspensions were placed on wheat florets. Following inoculation, the spikes were covered with polyethene bags for 72 h to maintain high humidity. The plants were subsequently maintained under glasshouse conditions at approximately 25°C during the day and 19°C at night. Five biological replicates were maintained per treatment.

Disease symptoms on wheat spikes and harvested seeds were recorded photographically. Histopathological examination of inoculated spike tissues was conducted to examine fungal structures associated with infection, including the fruiting bodies (Perithecia) observed in the examined tissues. To assess Koch’s postulates, symptomatic wheat tissues were used for fungal re-isolation, and the recovered isolates were compared with the original cultures.

### 2.4 Whole genome sequencing and analysis

Approximately 100 ng of genomic DNA of each isolate was enzymatically fragmented to get an average insert size of 200–300 bp for sequencing. Next, the fragmented DNA underwent end repair, A-tailing, and adapter ligation following the manufacturer’s instructions. Ligated fragments were purified using AMPure XP beads from Beckman Coulter and enriched by limited-cycle PCR using indexed primers. Final libraries were quantified using a Qubit fluorometer and assessed for fragment-size distribution using an Agilent 4200 TapeStation system with a D1000 ScreenTape.

Illumina libraries were normalised based on Qubit concentration and fragment size and sequenced as paired-end reads (2 × 150 bp) on an Illumina NovaSeq 6000 platform following the manufacturer’s protocol. Long-read sequencing libraries were prepared from approximately 800 ng of high-molecular-weight DNA using the Oxford Nanopore Technologies Ligation Sequencing Kit with Native Barcoding (SQK-LSK109 and EXP-NBD104). Barcoded libraries were pooled in equimolar concentrations and sequenced on an Oxford Nanopore PromethION P2 Solo flow cell.

### 2.5 Read processing and genome assembly

Raw Illumina reads were filtered using Trimmomatic v0.39 to remove adapter sequences, ambiguous bases (N > 5 %), and low-quality reads (SLIDINGWINDOW:10:25, LEADING:25, TRAILING:25, MINLEN:100). Quality assessment of Illumina reads was performed using FastQC and summarised using MultiQC. Nanopore reads were quality-filtered using NanoFilt, and sequencing statistics were generated using NanoPlot.

Hybrid de novo genome assembly was performed using MaSuRCA v4.1.4, followed by scaffolding with RagTag. The assemblies were polished using Medaka for Nanopore reads and Pilon for Illumina reads (2–3 rounds of polishing). Assembly quality was evaluated using QUAST, and Genome completeness was assessed using BUSCO with the *Hypocreales_odb10* lineage dataset (Fig. S1). Foreign contamination was screened and removed using the NCBI Foreign Contamination Screen (FCS) tool suite on the Galaxy platform (https://usegalaxy.org/). The cleaned assembly was then deposited in the DDBJ/ENA/GenBank database. The raw Illumina NovaSeq paired-end and Nanopore reads were deposited in the NCBI Sequence Read Archives (SRA) database.

### 2.6 Gene structure and functional annotation

Protein-coding genes in the newly sequenced Indian *Fusarium* isolates, *Fusarium graminearum* TNW1 and *Fusarium avenaceum* DMW8, were predicted using AUGUSTUS with the *Fusarium* model (Stanke et al. 2004). For comparative analysis, publicly available genome assemblies were retrieved from the National Center for Biotechnology Information (NCBI). Existing protein FASTA files were used for reference genomes where available. For reference assemblies lacking the required protein FASTA files, protein-coding genes were predicted using AUGUSTUS with the *Fusarium* model. The resulting gene models were processed using gffread to extract predicted protein-coding sequences for downstream comparative analyses.

Genome assemblies were renamed using a standardised nomenclature comprising the first three letters of the genus name, the first two letters of the species name and the isolate identifier (e.g., Fusas_140001 for *Fusarium asiaticum* isolate 140001). Assembly accession numbers were matched with isolate metadata, including strain, host and geographic origin, to generate a genome-renaming table. Genome completeness was assessed using BUSCO v6.1.0 (Tegenfeldt et al. 2025) with the Hypocreales odb10 lineage dataset comprising 4,494 conserved orthologous groups. Assembly and predicted protein completeness were evaluated using the numbers of complete, fragmented, and missing BUSCOs. Genomes exhibiting low completeness or substantial fragmentation were excluded from downstream analyses.

Comparative genome selection considered BUSCO completeness, assembly fragmentation, host association, and geographic representation. Wheat-associated isolates were prioritised; when unavailable, genomes from other suitable hosts were considered. Where multiple genomes originated from the same geographic location, preference was given to assemblies with higher BUSCO completeness and lower assembly fragmentation. Initially, 150 publicly available genome assemblies of *F. graminearum* and 15 of *F. avenaceum* were considered alongside the newly sequenced isolates. Following quality assessment and filtering, 21 *F. graminearum* genomes and 15 *F. avenaceum* genomes were retained, yielding 36 genomes for comparative analysis. Genome accession numbers, isolate metadata, and quality statistics are provided in Supplementary File S2. The NCBI database was accessed on 8 October 2026.

Functional annotation was performed using OmicsBox with DIAMOND BLASTP, eggNOG v2.1.12 (Cantalapiedra et al., 2021), Phobius (Käll et al., 2007) and InterProScan (Jones et al., 2014). Annotations were integrated using the Funannotate annotate process. BLAST-derived accession identifiers were used to retrieve gene names and UniProt identifiers from PIR, UniProt/Swiss-Prot, TrEMBL, RefSeq, GenPept, UniGene and PDB databases for Gene Ontology (GO) mapping, annotation, GO-Slim classification and functional enrichment analysis. A standardised annotation application was also developed to support reproducible protein annotation.

To perform KEGG pathway annotation, Protein FASTA sequences were first submitted to the KEGG Automatic Annotation Server (KAAS; https://www.genome.jp/kegg/kaas/) to obtain gene-to-KEGG Orthology (KO) assignments, which were provided through a KAAS result URL (Moriya et al., 2007). As the analysis involved 36 *Fusarium* genomes, manually retrieving and processing the KAAS KO-assignment results was cumbersome. Therefore, we developed a local Python/Streamlit-based KAAS–KEGG Pathway Analyser, with the source code made available through GitHub. The completed KAAS result URL containing the KO assignments was provided as input to the application, which automated the processing of KO numbers, the retrieval of functional, EC, and KEGG pathway annotations, protein-level mapping, and the generation of integrated Excel output for downstream comparative analysis. Biopython v1.87 (Chapman and Chang, 2000) was used for data processing. Additionally, to investigate pathogenicity- and virulence-related functions, a curated subset of KEGG pathways was selected, encompassing core signalling, host interaction, and secondary metabolism processes relevant to fungal pathogenicity. These included key signalling pathways (MAPK, Ras, and cAMP) known to regulate fungal development, stress adaptation, morphogenesis, and infection processes (Ma et al., 2013; Kazan et al., 2012).

The Telomere Identification Toolkit (tidk) v0.2.65 was applied to the selected genomes to identify and compare telomeric motifs. Using a 1,000-bp sliding window, the standard fungal telomeric pattern TTAGGG was used to detect telomeres in fungi (Fulnečková et al. 2013).

### 2.7 Identification of non-coding RNA genes (rRNA and tRNA)

Ribosomal RNA (rRNA) genes were identified in the selected Fusarium genome assemblies using “Barrnap v0.9” (Kseemann, 2014) with the eukaryotic model. Barrnap was run with the default settings, and the FASTA and GFF formats of the predicted rRNA sequences were obtained for further examination*. “tRNAscan-SE v2.0.12”* (Chan et al., 2021). was used to predict transfer RNA (tRNA) genes in eukaryotic mode. Covariance models implemented in Infernal were used to identify tRNAs in genome assemblies supplied as input. Genomic coordinates were translated to BED format, and the data were produced in tabular form. "Bedtools" and "getfasta" were used to retrieve strand-corrected tRNA sequences. The total number of expected tRNA and rRNA genes for each genome was determined and utilised for cross-species comparison.

### 2.8 Secretome analysis and pathogenicity-related genes using a computational pipeline

Predicted protein sequences from the selected genomes were analysed for secretion using SignalP v6.0 (Teufel et al. 2022), and proteins lacking signal peptides were excluded. The remaining proteins were filtered using DeepTMHMM v1.0 (Hallgren et al. 2022) to remove sequences containing transmembrane helices, followed by NetGPI v1.1 (Gíslason et al. 2021) to exclude GPI-anchored proteins. Subcellular localisation was predicted using TargetP v2.0 (Armenteros et al. 2019) and DeepLoc v2.1 (Ødum et al. 2024), and only proteins predicted to be extracellular were retained. The resulting secretome was used to predict effector proteins using Effector v3.0 (Sperschneider and Dodds 2022). (Fig S2) The relative abundance of cysteine (C), serine (S), and tryptophan (W) residues was calculated as a percentage of protein length using custom Python scripts, given their established roles in fungal effector structure and function. Small cysteine-rich (SCR) effectors were defined as predicted effector proteins with a length ≤300 amino acids and a cysteine content ≥4%, restricted to secreted extracellular proteins. Identified candidate secreted effector proteins (CSEPs) were further subjected to Gene Ontology (GO) analysis using Blast2GO, and functional characterisation was performed using InterProScan and the PHI-base database to identify domains, protein families. Fig S2.

### Prediction of secondary metabolite gene clusters

Secondary metabolite biosynthetic gene clusters (BGCs) were predicted using antiSMASH fungal v7.1.0 (Blin et al., 2023). The selected *Fusarium* genome assemblies were analysed with default antiSMASH parameters, and secondary metabolite regions were identified with relaxed strictness. The resulting biosynthetic gene clusters were manually examined and verified using BLASTP against the NCBI database.

### Identification and annotation of carbohydrate active enzyme classes

The predicted protein datasets from the selected *Fusarium* genomes were analysed using the dbCAN3 server (Zheng et al., 2023) via HMMER searches against the dbCAN database (E-value < 1 × 10 ¹ ; coverage > 0.35). The identified carbohydrate-active enzymes (CAZymes) were classified into the six major CAZy classes: glycoside hydrolases (GHs), glycosyltransferases (GTs), auxiliary activities (AAs), carbohydrate esterases (CEs), polysaccharide lyases (PLs), and carbohydrate-binding modules (CBMs) (Hao et al., 2021).

### 2.9 Analysis of Transposable element annotation, TE insertions and repeat-induced point mutation (RIP)

Transposable elements (TEs) were annotated in the selected *Fusarium* genomes using the Earl Grey pipeline (v7.2.1). The Earl Grey pipeline integrates both de novo and homology-based approaches for comprehensive TE annotation. De novo TE families were identified using RepeatModeler v2.0.7 (Baril et al., 2024), followed by iterative consensus refinement using the ‘*BLAST–Extend–Align–Trim’* (BEAT) algorithm (5 iterations, up to 20 sequences per family).

RepeatMasker was configured to use RMBlast (v2.14.1+) as the primary sequence-similarity search engine. In contrast, HMMER v3.4 (Wheeler & Eddy, 2013) was used internally for profile hidden Markov model (HMM)-based searches against ‘*Dfam’* (Hubley et al., 2016). This hybrid approach enables the detection of both conserved and highly divergent TE families. An initial step using known fungal repeats was performed before de novo repeat discovery, and the resulting de novo libraries were integrated with curated databases for final genome-wide annotation.

Long terminal repeat (LTR) retrotransposons were identified using LTR_Finder v1.07 (Xu & Wang, 2007), while tandem and simple repeats were detected using Tandem Repeats Finder (TRF) and additional repeat-detection tools integrated within the pipeline. TE annotations were further refined using ‘*RepeatCraft*’ to merge fragmented elements and resolve nested insertions. Kimura two-parameter (K2P) divergence values were calculated to estimate the relative age of TE insertions and to generate repeat landscape profiles. All genomes were processed using four CPU threads. Final outputs included TE classification summaries, genome coverage statistics, curated TE consensus libraries, genome-wide TE annotations in GFF format, and TE landscape analyses.

### Repeat-induced point mutation (RIP) analysis

Repeat-induced point mutation (RIP) activity was assessed using a sliding-window approach following the methodology of TheRIPper v3.0 (van Wyk et al., 2019). The selected genomes were scanned using 1,000-bp windows with a 500-bp step size. The RIP composite index was calculated from dinucleotide frequencies; windows with values greater than 0 were considered RIP-affected, and values were capped at 5.

Consecutive RIP-positive windows were merged to define large RIP-affected regions (LRARs), requiring at least seven consecutive windows. Transposable elements (TEs) longer than 100 bp were intersected with LRARs using BEDTools v2.32.1 (Quinlan & Hall, 2010), with a minimum overlap of 50%. Genome-wide RIP burden was calculated as the proportion of genomic bases contained within LRARs and visualised in R v2026.01.1+403 (R Core Team, 2026).

### Detection of TE insertions in newly sequenced Indian Fusarium strains

To investigate transposable element (TE) proliferation at an intraspecific level, we analysed TE insertions in two *Fusarium* strains, DMW8 and TNW1, using their respective assembled reference genomes generated in this study. Paired-end Illumina sequencing reads were assessed for quality using FastQC v.0.11 (Andrews 2010) and subsequently trimmed with fastp v.0.23.4 using default parameters and a minimum read length of 60 bp (Chen et al. 2018). Sequencing reads were aligned to their respective reference genomes using BWA (Li and Durbin 2009) as implemented within the McClintock pipeline v2.0 (Chen et al. 2023).

TE insertions were detected using multiple approaches integrated into McClintock (Nelson et al., 2017), including ngs_te_mapper2 (Han et al., 2021), RelocaTE (Robb et al., 2013), RelocaTE2 (Chen et al., 2017), TEMP (Zhuang et al., 2014), TEMP2 (Yu et al., 2021), RetroSeq (Keane et al., 2013), PoPoolationTE2 (Kofler et al., 2016), TE-locate (Platzer et al., 2012) and TEFLoN (Adrion et al., 2017). Following a comparative evaluation of the outputs generated by these methods, the results from TEFLoN were selected for downstream analyses due to their robustness and consistency.

Consensus sequences of TE families generated by the ‘*Earl Grey pipeline’* were supplied to McClintock using the *‘-c option’*, and genomic locations of TE families were incorporated using the *‘-g option’* when available. McClintock was used to identify both reference and non-reference TE insertions across each genome. The number of TE insertions per megabase (Mbp) and their distribution among TE orders and superfamilies were quantified (Wickham, 2016).

### 3. Comparative orthology and Phylogenomics analysis

To provide evolutionary and comparative genomic context, OrthoFinder v2.5.5 was used to cluster the predicted protein sequences of the 36 selected *Fusarium* genomes into orthogroups and infer a genome-scale species tree (Emms and Kelly, 2019). The coalescent-based species tree was generated using the STAG (Species Tree Inference from All Genes) algorithm and rooted with STRIDE as implemented in OrthoFinder (Emms and Kelly, 2018). The resulting orthogroups and gene-count matrix were used for comparative gene-family analysis.

A complementary concatenation-based phylogenomic analysis was performed using single-copy orthologous genes shared across the selected taxa. Protein sequences were aligned using MAFFT (Katoh and Standley, 2013), trimmed with trimAl v1.4.rev15 (Capella-Gutiérrez et al., 2009), and concatenated using AMAS (Borowiec, 2016). The concatenated dataset was used to infer a maximum-likelihood phylogeny using RAxML-NG (Kozlov et al., 2019), with genes partitioned under the JTT + G4 amino acid substitution model, and branch support was assessed by bootstrap analysis.

Subsequently, a separate set of 10 genomes of various FHB-associated species, including the two newly sequenced isolates *F. graminearum* TNW1 and *F. avenaceum* DMW8, along with eight reference genomes, was selected for focused evolutionary and comparative analysis. The predicted protein datasets of TNW1 and DMW8 generated in this study were compared with the corresponding reference genomes *F. graminearum* strain PH-1 (GCF_ 000240135.3) and *F. avenaceum* strain FaLH03 (GCA_ 000769305.1), respectively. The remaining reference genomes included *F. culmorum* strain Class2-1B (GCA_016952355.1), *F. poae* strain DAOMC 252244 (GCF_019609905.1), *F. asiaticum* strain KCTC 16664 (GCA_025258505.1), and *F. boothii* strain CBS 316.73 (GCA_017656985.1). Additionally, *F. graminearum* strain mFG23 (GCA_051903665.1) and *F. avenaceum* strain WV21P1A (GCA_025948275.1) were included as additional representatives of the respective species because of their high genome completeness and assembly quality, providing suitable wheat-associated reference genomes for comparison with the newly sequenced isolates.

The same 10 genomes were also analysed using OrthoVenn3 (Sun et al., 2023) via a locally deployed Docker instance to perform orthologous gene cluster comparisons and annotations. The 36-genome analysis was not feasible in OrthoVenn3 because of its computational/genome-number limitations.

CAFE5 (Mendes et al., 2021) was used to estimate gene-family expansions and contractions for both the 36-genome and 10-genome datasets. For the 10-genome CAFE5 analysis, OrthoFinder-derived gene counts and the corresponding species tree were used. Species trees were rooted and converted to ultrametric trees before CAFE5 analysis, and significant expansions or contractions were identified at P ≤ 0.05.

Pathogenicity-related clusters were identified by screening standardised functional annotations against predefined keywords that represent major virulence-associated pathways, including MAPK–HOG/CWI, cAMP, TOR, calcium–calcineurin signalling, cell wall-degrading enzymes (CWDEs), secondary metabolism, and core virulence factors. Clusters containing at least one predefined keyword were classified as pathogenicity-related (Kazan et al., 2012; Ma et al., 2013; Rauwane et al., 2020).

In addition to phylogenetic reconstruction, comparative genomic metrics, including assembly N50, genome size, number of predicted gene models, and the number of unassigned genes identified by OrthoFinder, were evaluated to facilitate interpretation of genomic diversity among species.

## 4. Results

### 4.1 Morphological and molecular identification of FHB-associated Fusarium isolates

Morphological characterisation revealed that *Fusarium graminearum*, TNW1, shows a large amount of dense aerial mycelium, ranging from Cottony to fluffy. The colony shows a Carmine red colour. Red-brown to orange sporodochia are produced slowly (> 30 days) in PDA and in Carnation Leaf Agar (CLA), along with thick-walled, moderately curved macroconidia with 5–6 septa. Two types of macroconidia are produced: primary and secondary macroconidia. Primary macroconidia: 1-3 septate, 15-25μm × 3.75-5 μm produced from a conidiogenous cell. Secondary macroconidia: 4-7 septate, narrow, slender, long, straight to curved, 40-47.5 µm × 3.7-5 µm. Apical cell: elongated, and basal cell: foot-shaped.

The colonies of *F. avenaceum* DMW8 were typically characterised by loosely fluffy to cottony aerial mycelia fringed with white and pink to carmine pigmentation. Macroconidia from sporodochia were uniform, 5-7-septate, narrowly fusoid, curved, with an elongated apical cell and a well-marked foot cell. The size of 5-septate macroconidia was 63.8 (51.1-80.0) µm × 4.1 (3.6-4.7) µm, with an apical cell length of 17.1µm (14.1-21.2 µm). In the aerial mycelium, fusoid 0-3-septate conidia (10-26 × 2.2-4.2 µm) occurred occasionally (Fig. 1a).

**Fig. 1:**
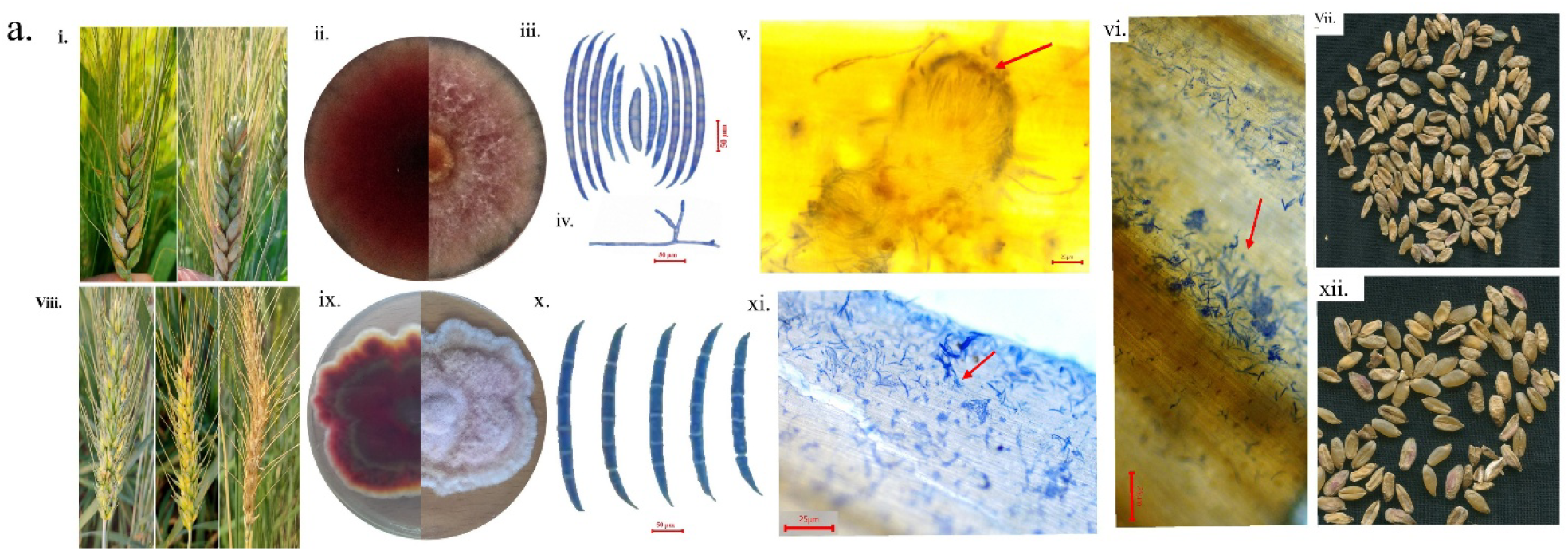
Morphological and pathological characteristics of wheat Fusarium head blight (FHB)-associated isolates, *Fusarium graminearum* TNW1 and *Fusarium avenaceum* DMW8. (i–vii) Characteristics associated with *F. graminearum* TNW1, showing (i) wheat spikes exhibiting floret bleaching and tombstone kernel symptoms, (ii) colony morphology and pigmentation on potato dextrose agar (PDA; 7 days), (iii) macroconidial morphology (scale bar = 50 µm), (iv) conidiophore morphology (scale bar = 50 µm), (v) perithecia observed in infected wheat spike tissue (scale bar = 25 µm), (vi) fungal structures observed on infected wheat spikelet tissue (scale bar = 25 µm), and (vii) seeds harvested from infected wheat spikes showing tombstone kernels. (viii–xii) Characteristics associated with *F. avenaceum* DMW8, showing (viii) wheat spikes exhibiting FHB symptoms, (ix) colony morphology and pigmentation on PDA (7 days), (x) macroconidial morphology (scale bar = 50 µm), (xi) fungal structures observed on infected wheat spikelet tissue (scale bar = 25 µm), and (xii) seeds harvested from infected wheat spikes showing tombstone kernels.

Based on morphological similarity and multi-locus phylogeny with high sequence similarity (≥99%) to reference strains in GenBank, isolate TNW1 was identified as *F. graminearum* Schwabe. In contrast, DMW8 was identified as *F. avenaceum* (Fr.). *Sacc*. Further Phylogenetic analysis confirmed that isolates DMW8 and TNW1 cluster with reference strains of *F. avenaceum* and *F. graminearum*, respectively (Fig. 1b). This identity was again confirmed at the ICAR–National Bureau of Agriculturally Important Microorganisms (NBAIM), Uttar Pradesh, India. Pure cultures were deposited at NBAIM under accession numbers NAIMCC-F-04729 (TNW1) and NAIMCC-F-04728 (DMW8). The barcode sequences were deposited in NCBI under accession numbers PX567756 and PX567757 (ITS), PX578949 and PX578950 (TEF1-α), and PZ015365 and PZ015366 (RPB2), and are available in the public domain.

**Fig. 1b.**
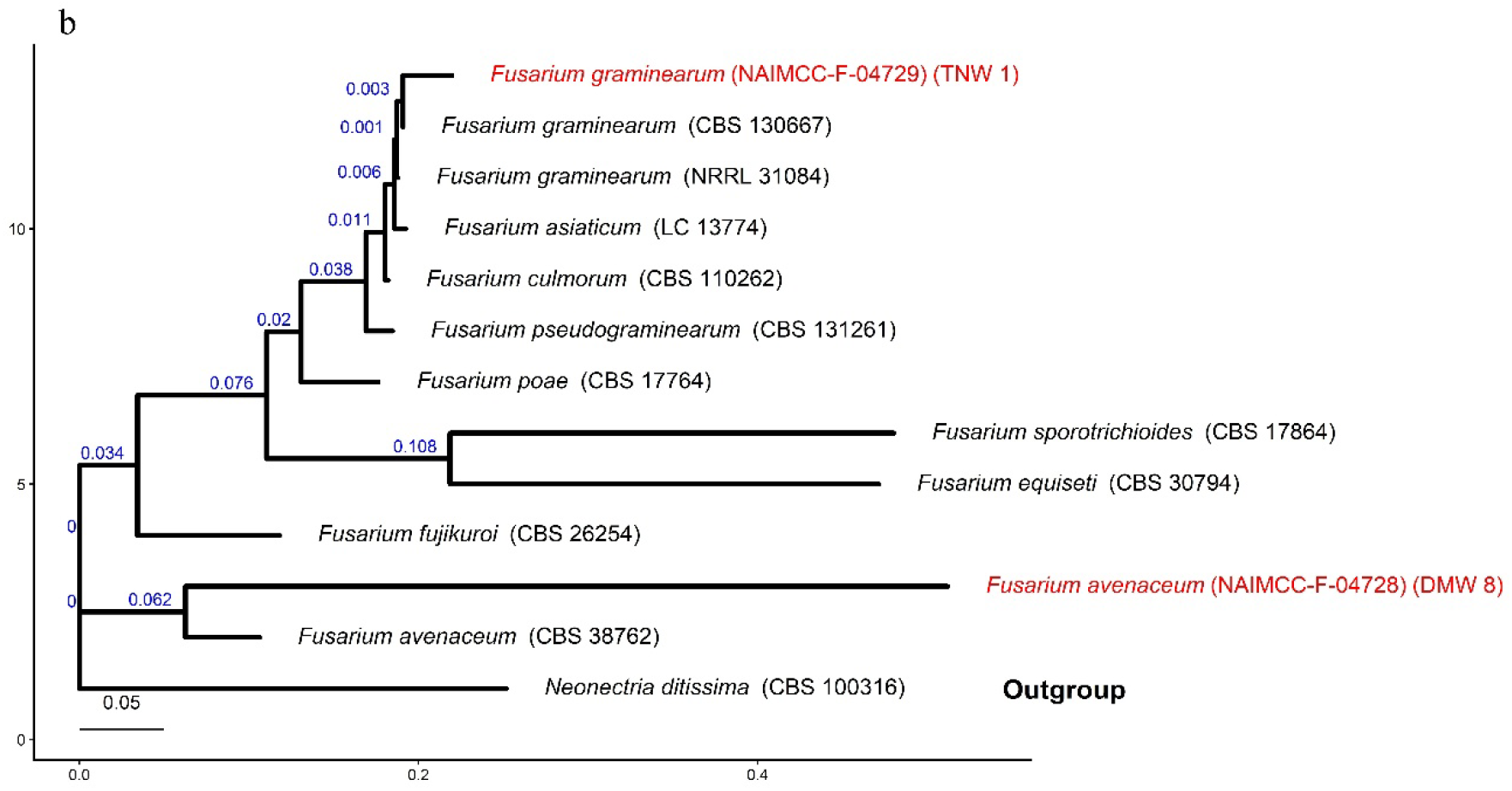
Maximum-likelihood phylogenetic tree based on concatenated ITS, TEF1-α, and RPB2 sequences showing the relationships of the studied isolates with reference Fusarium species. Bootstrap values (≥70%) based on 1000 ultrafast replicates are shown at the nodes. Isolates obtained in this study are indicated in red. Neonectria ditissima (CBS 100316) was used as the outgroup. The scale bar represents substitutions per site.

### 4.2 Pathogenicity Symptoms and Histopathological Observations

Inoculation with *Fusarium graminearum* TNW1 and *Fusarium avenaceum* DMW8 was associated with visible disease symptoms on wheat spikes and harvested seeds (Fig. 1). Histopathological examination of inoculated spike tissues revealed fungal fruiting bodies (Fig. 1), providing additional microscopic evidence of fungal development in the affected tissues.

The observed spike and seed symptoms, together with the histopathological findings, provided phenotypic evidence complementary to the genomic characterisation of TNW1 and DMW8. The observed symptoms on wheat spikes and seeds, together with the histopathological evidence of fungal structures, supported the pathogenic potential of *F. graminearum* TNW1 and *F. avenaceum* DMW8 on wheat. Re-isolated fungi showed morphological characteristics consistent with the original isolates, supporting their association with the observed disease symptoms (Fig. 1a).

### 4.3 Genome assembly characteristics and completeness of Indian Fusarium isolates

Hybrid genome sequencing of the Indian isolates *Fusarium graminearum* TNW1 (GCA_054643435.1) and *Fusarium avenaceum* DMW8 (GCA_054643395.2) generated draft genome assemblies of 36.29 Mb and 42.00 Mb, respectively (Table 2). The TNW1 assembly comprised 52 contigs and 21 scaffolds, whereas DMW8 comprised 217 contigs and 35 scaffolds. The largest contigs measured 10,031,799 bp in TNW1 and 6,221,714 bp in DMW8. The corresponding N50 values were 4,271,658 bp and 4,219,374 bp, with auN values of 4,942,144 bp and 4,743,021 bp and L50 values of 3 and 4, respectively. GC content was 48.27% in TNW1 and 48.58% in DMW8. Sequencing coverage depths were 162.6× and 214.3×, respectively, with 99.99% of each genome covered at a minimum depth of 1×.

**Table 2.** Comparative summary of genome assembly metrics, completeness assessment (BUSCO; Hypocreales odb10), and functional annotation features for newly sequenced strains *Fusarium graminearum* TNW1 and *Fusarium avenaceum* DMW8

| **Feature / Category** | **F. graminearum (TNW1)** | **F. avenaceum (DMW8)** |
|---|---|---|
| **NCBI Accession number** | **GCA_054643435.1** | **GCA_054643395.2** |
| **Assembly Statistics** |  |  |
| **Estimated genome size (Mb)** | 36.29 | 42 |
| **Number of contigs** | 52 | 217 |
| **Number of scaffolds** | 21 | 35 |
| **Largest contig (bp)** | 10031799 | 6221714 |
| **GC content (%)** | 48.27 | 48.58 |
| **N50 (bp)** | 4271658 | 4219374 |
| **auN (bp)** | 4942144 | 4743021 |
| **L50** | 3 | 4 |
| **Average coverage depth (×)** | 162.6× | 214.3× |
| **Genome coverage ≥1× (%)** | 99.99 | 99.99 |
| **BUSCO Analysis (Hypocreales_odb10)** |  |  |
| **Total BUSCO groups searched** | 4494 | 4494 |
| **Complete BUSCOs (C)** | 4463 (99.3%) | 4386 (97.6%) |
| **Complete & single-copy BUSCOs (S)** | 4463 (99.3%) | 4381 (97.5%) |
| **Complete & duplicated BUSCOs (D)** | 0 (0%) | 5 (0.1%) |
| **Fragmented BUSCOs (F)** | 4 (0.1%) | 17 (0.4%) |
| **Missing BUSCOs (M)** | 27 (0.6%) | 91 (2.0%) |
| **Annotation Statistics** |  |  |
| **Protein-coding genes** | 11750 | 13028 |
| **mRNAs** | 11750 | 13028 |
| **tRNAs** | 320 | 302 |
| **rRNAs** | 69 | 60 |
| **Average gene length (bp)** | 1709.47 | 1693 |
| **Average CDS length (bp)** | 534.6 | 528.3 |
| **Number of exons** | 34153 | 32663 |
| **Average exon length (bp)** | 534.6 | 528.3 |
| **Average protein length (aa)** | 516.96 | 510.83 |
| **Proteins with InterProScan annotation** | 10837 | 10406 |
| **Proteins with eggNOG annotation** | 11366 | 12777 |
| **Proteins with Pfam annotation** | 8921 | 8642 |
| **Proteins with MEROPS annotation** | 334 | 315 |
| **Secretory proteins** | 1235 | 1110 |

Assembly completeness was assessed using BUSCO against 4,494 conserved orthologous groups from Hypocreales_odb10 (Table 2). TNW1 contained 4,463 complete BUSCOs (99.3%), all of which were single copy, with no duplicated BUSCOs. Four BUSCOs (0.1%) were fragmented, while 27 (0.6%) were missing. DMW8 contained 4,386 complete BUSCOs (97.6%), including 4,381 single-copy (97.5%) and five duplicated (0.1%) BUSCOs. Seventeen BUSCOs (0.4%) were fragmented, and 91 (2.0%) were missing. Thus, TNW1 showed higher BUSCO completeness and fewer missing orthologs than DMW8. Genome assembly quality was further visualised using Circos diagrams (Figs. 2 and 3). The two assemblies were included in a comparative dataset of 36 genomes, comprising 21 *F. graminearum* and 15 *F. avenaceum* genomes selected following quality assessment and consideration of host association and geographic representation (Supplementary File S2).

**Fig. 2.**
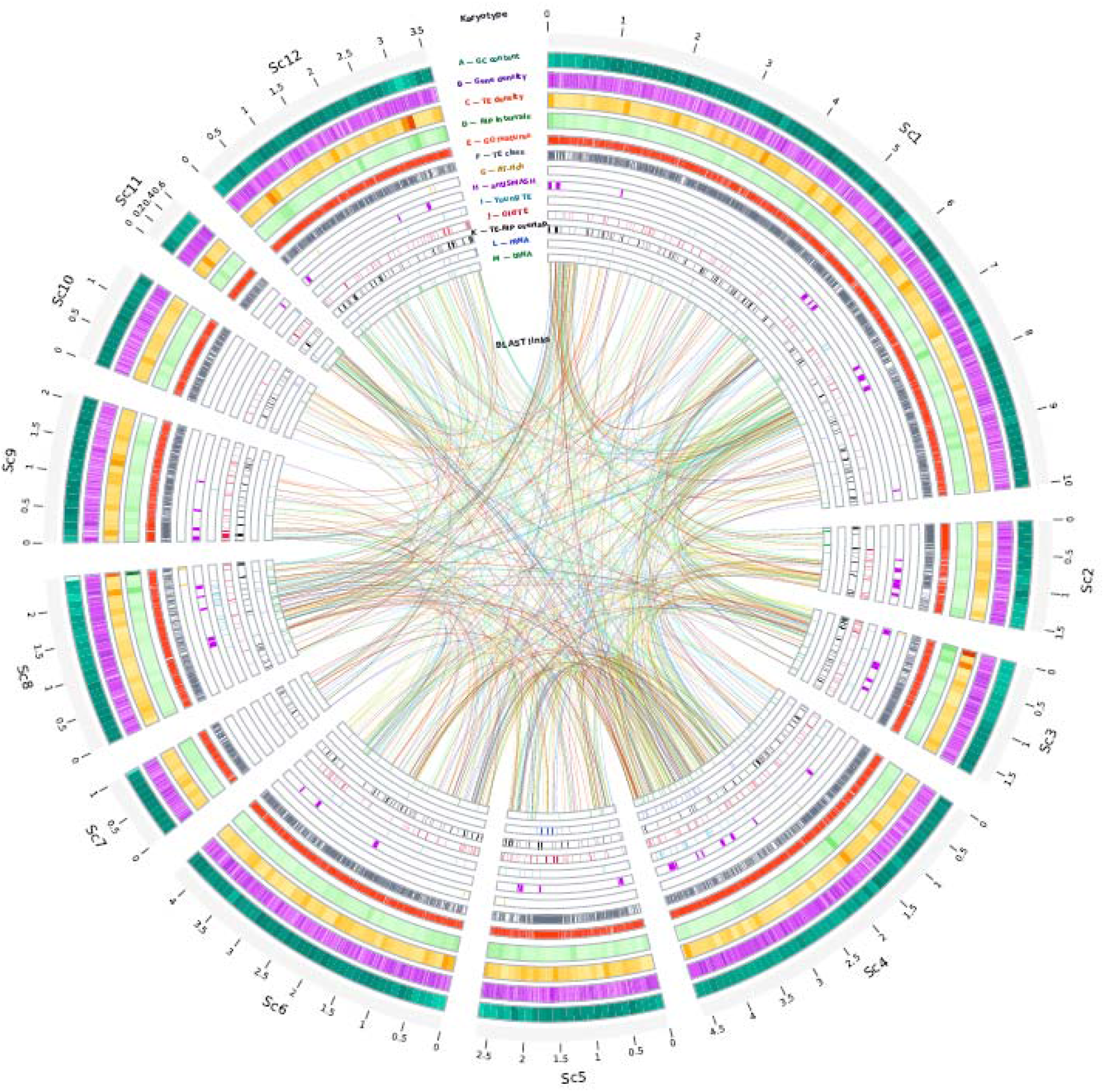
Circos plot of the *F. graminearum* TNW1 genome. The outer ring represents scaffolds (Sc1 – Sc12) and genomic positions in megabases (Mb). Tracks **A**: GC content, **B**: gene density, C: TE density, D: RIP intervals, E: GO features, F: TE class, G: AT-rich regions, H: antiSMASH regions, I: young TEs, J: old TEs, K: TE–RIP overlap, L: rRNA, M: tRNA. Central colored ribbons represent genomic links after filtering and subsampling. Scaffolds shorter than 100 kb (09 scaffolds) were excluded.

**Fig. 3.**
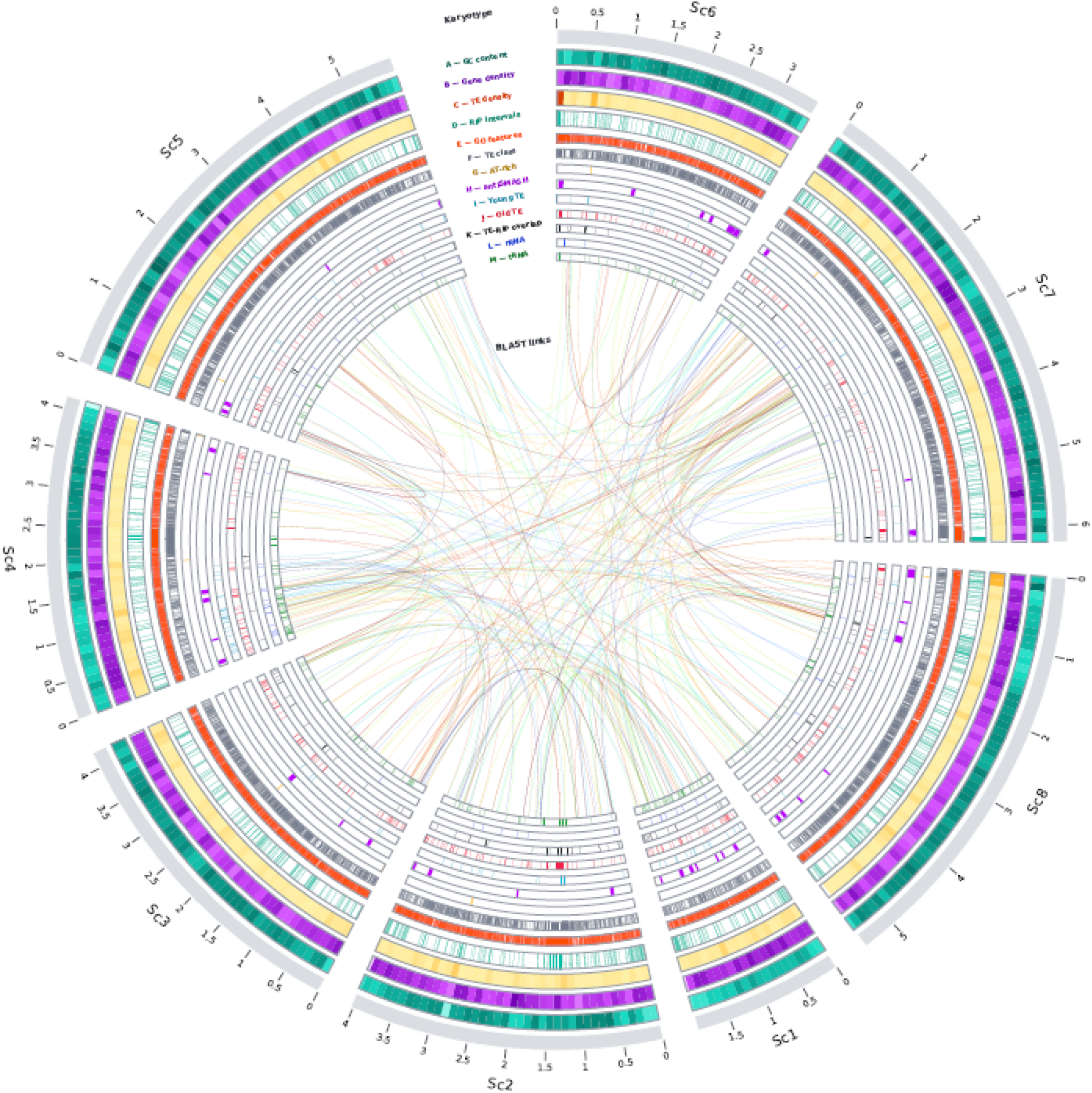
Circos plot of the *F. avenaceum,* DMW8 genome. The outer ring represents scaffolds (Sc1 – Sc8) and genomic positions in megabases (Mb). A: GC content, B: gene density, C: TE density, D: RIP intervals, E: GO features, F: TE class, G: AT-rich regions, H: antiSMASH regions, I: young TEs, J: old TEs, K:TE–RIP overlap, L: rRNA, M: tRNA. Central colored ribbons represent filtered and subsampled genomic links. Scaffolds shorter than 100 kb (27 scaffolds) were excluded.

### 4.4 Comparative functional annotation reveals differences in gene repertoires and pathway representation

Gene prediction and functional annotation were performed for the 36 selected *Fusarium* genomes to characterise their predicted gene repertoires and functional profiles (Table 2; Supplementary File S3). The *Fusarium graminearum* TNW1 genome contained 11,750 protein-coding genes, whereas *Fusarium avenaceum* DMW8 contained 13,028. Across the comparison genomes, protein-coding gene counts ranged from 11,328 to 14,132 in *F. graminearum* and from 11,233 to 15,886 in *F. avenaceum*. The average predicted gene lengths were 1,709.47 bp in TNW1 and 1,693.00 bp in DMW8, while the average protein lengths were 516.96 and 510.83 amino acids, respectively. TNW1 contained 34,153 exons, compared with 32,663 in DMW8. The predicted non-coding RNA complement comprised 320 tRNAs and 69 rRNAs in TNW1 and 302 tRNAs and 60 rRNAs in DMW8 (Table 2; Supplementary Table S3.1).

Functional annotation identified 11,366 eggNOG-annotated proteins in TNW1 and 12,777 in DMW8. InterProScan annotated 10,837 and 10,406 proteins, respectively, whereas Pfam annotation covered 8,921 proteins in TNW1 and 8,642 in DMW8. MEROPS identified 334 proteins in TNW1 and 315 in DMW8, while the predicted secretomes comprised 1,235 and 1,110 proteins, respectively (Table 2). Across the comparison genomes, eggNOG annotation counts ranged from 11,057 to 12,862 in *F. graminearum* and from 11,096 to 13,359 in *F. avenaceum*. InterProScan counts ranged from 10,732 to 12,423 and from 10,816 to 13,303, respectively, whereas Pfam counts ranged from 8,851 to 9,923 in *F. graminearum* and from 9,243 to 10,688 in *F. avenaceum*. MEROPS annotation counts ranged from 395 to 437 in *F. graminearum* and from 400 to 463 in *F. avenaceum*. Predicted secretome sizes ranged from 1,157 to 1,274 proteins in *F. graminearum* and from 1,120 to 1,456 proteins in *F. avenaceum* (Supplementary File S3). Overall, TNW1 and DMW8 exhibited differences in their predicted functional annotation profiles. In particular, both isolates had fewer MEROPS-annotated proteins than their corresponding comparison genomes, while DMW8 also showed lower counts of InterProScan and Pfam annotations than the other *F. avenaceum* genomes.

KEGG Orthology (KO) analysis assigned terms to 4,306 proteins in TNW1 and 4,251 in DMW8. In both genomes, metabolism was the largest functional category, followed by genetic information processing and cellular processes (Fig. 4a). The KEGG annotation proportions were 36.65% for TNW1 and 32.63% for DMW8 (Supplementary Table S3.10). These results indicate broadly similar distributions of major functional categories, alongside differences in the proportion of genes assigned KO terms.

**Fig. 4.**
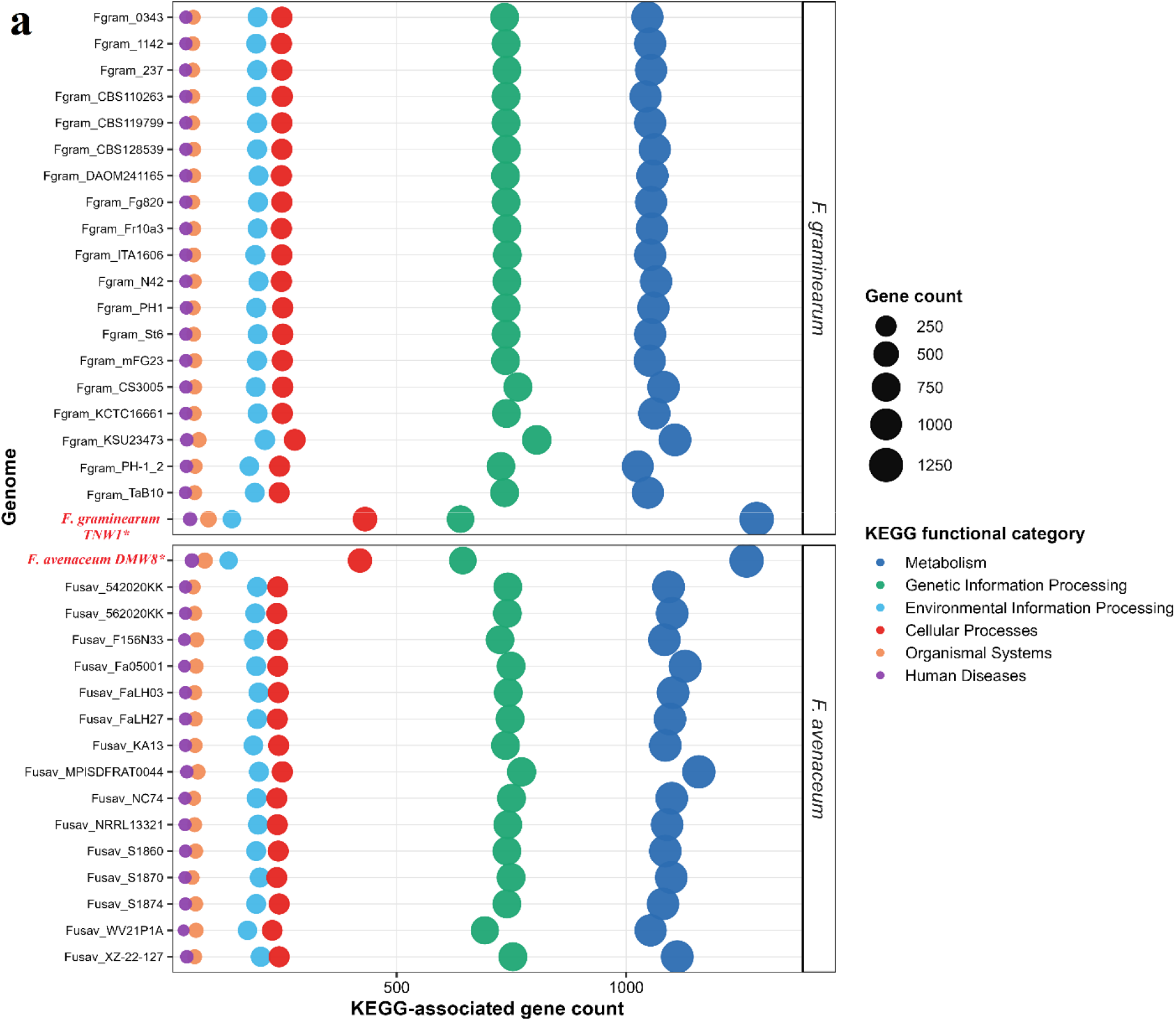
(a) Distribution of KEGG Level-1 functional categories across the analysed Fusarium genomes, showing the number of annotated genes assigned to each functional category.

Analysis of selected KEGG pathways revealed differences in the representation of specific functional groups (Fig. 4b; Supplementary Table S3.12). Among the analysed *F. graminearum* genomes, TNW1 had 19 genes assigned to terpenoid backbone biosynthesis, compared with 15–17 in the comparison genomes, and 16 genes associated with SNARE interactions in vesicular transport, compared with 13–15 in the other isolates. TNW1 also contained 11 genes in the MAPK signalling pathway, 23 in ABC transporters, two in cAMP signalling and five in Ras signalling, while Rap1 signalling was represented by three genes. In DMW8, terpenoid backbone biosynthesis and the FoxO signalling pathway were represented by 17 and eight genes, respectively. The remaining selected pathways showed variation in gene counts relative to the comparison genomes (Fig. 4b; Supplementary File S3). Overall, the annotation results demonstrate differences in predicted gene numbers, annotation coverage, and selected KEGG pathway assignments among TNW1, DMW8, and their respective comparison genomes.

**Fig. 4.**
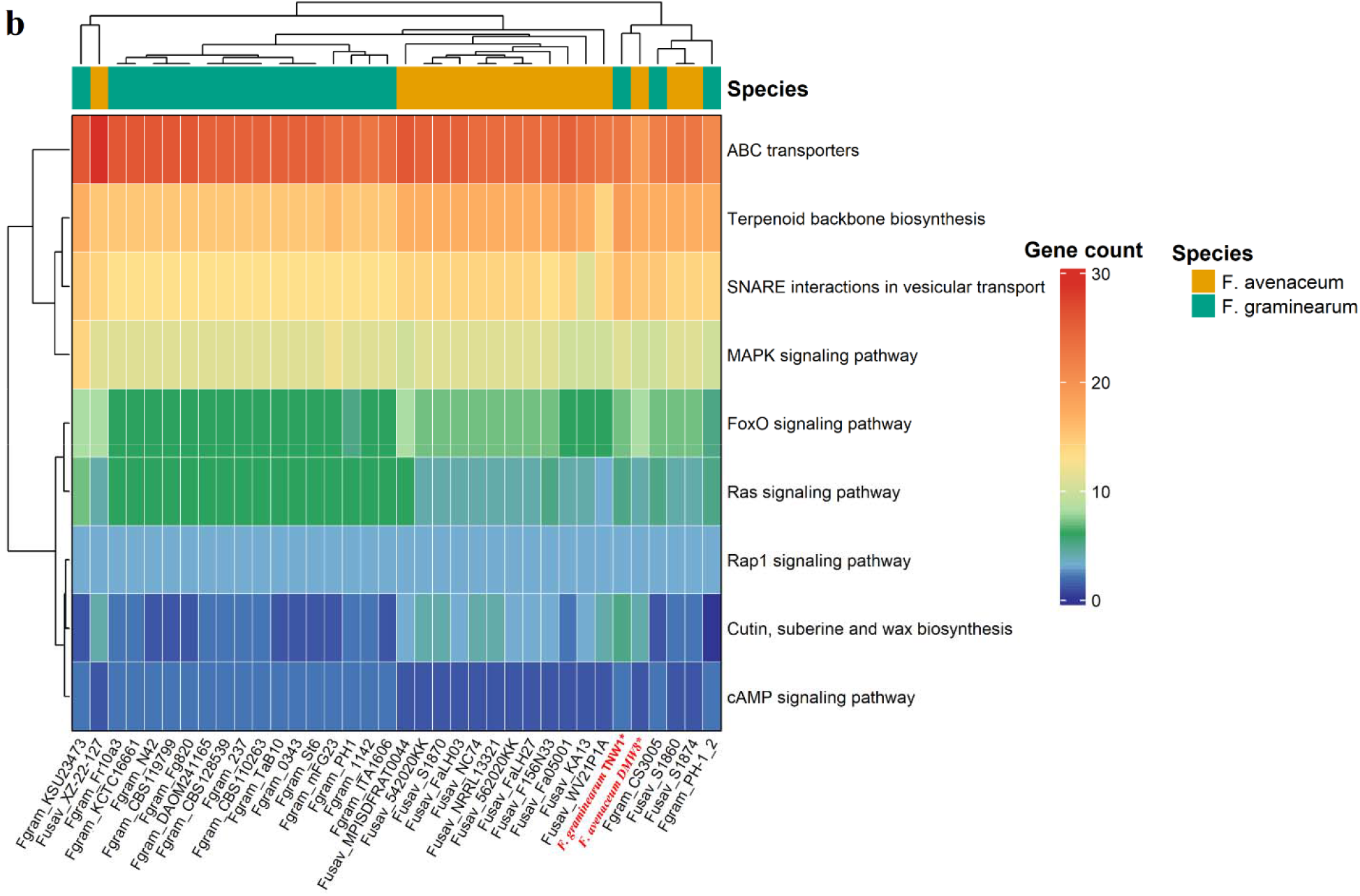
(b) Heatmap showing selected KEGG pathways associated with virulence, signaling, and secondary metabolism across Fusarium species and the isolates F. graminearum TNW1 and F. avenaceum DMW8. Colors represent log□-transformed gene counts.

### 4.5 Telomeric repeat profiles fall within the ranges observed across species-specific genome sets

TNW1 contained 5,771 TTAGGG repeat-containing 1-kb windows and 6,450 total repeat counts, within the ranges observed across the 21 *F. graminearum* genomes (5,651–6,585 windows and 6,280–7,600 total repeat counts). DMW8 contained 7,615 repeat-containing windows and 8,584 total repeat counts, within the ranges observed across the 15 *F. avenaceum* genomes (7,573–9,537 windows and 8,575–11,812 total repeat counts). Mean repeat counts were 0.178 in TNW1 and 0.204 in DMW8, with maximum counts of 17 and 26, respectively. Overall, both genomes showed TTAGGG repeat profiles within the ranges observed for their respective reference genome sets. (Supplementary file S4)

### 4.6 Distinct effector repertoires and species-associated secondary-metabolite biosynthetic profiles

Secretome analysis identified 1,235 (10.51%) secreted proteins in TNW1, including 325 predicted effector candidates. These effectors exhibited typical fungal virulence characteristics, with a mean protein length of 261.6 amino acids and an enrichment of cysteine residues; 85 proteins were classified as small cysteine-rich effectors (Fig. S5). Functional annotations revealed enrichment of extracellular enzymes and virulence-associated proteins, including carbohydrate-active enzymes, LysM domain proteins, proteases, and other cell wall-degrading enzymes (Fig. 5). In DMW8, 1,110 (8.52%) proteins were predicted to be secreted, including 285 effector candidates. These proteins had a mean length of 232.8 amino acids, and 92 were identified as small cysteine-rich effectors. Functional profiling indicated enrichment in glycoside hydrolases, cutinases, lectin/glucanase proteins, hydrophobins, and necrosis-inducing proteins (Fig. 5), supporting roles in host colonisation and extracellular interaction (supplementary file S5). Comparative analysis of predicted secondary metabolite biosynthetic gene clusters (BGCs) revealed conserved and variable profiles within the two *Fusarium* species. Conserved clusters included aurofusarin, chrysogine, clavaric acid, koraiol, orcinol/orsellinic acid, and oxyjavanicin. *F. graminearum* TNW1 additionally contained zearalenone, fusarielin H, fusaristatin A, and DON-related clusters, whereas DMW8 (*F. avenaceum*) contained ilicicolin H/J, fosfonochlorin, beauvericin, and hexadehydroastechrome clusters. DMW8 contained 16 known clusters, all detected in at least one reference genome, with up to 15 shared clusters, whereas TNW1 contained 18 clusters, of which 17 were shared with at least one reference genome; the hydroxyl-Fumonisin C1/Fumonisin C1/C2/C3/B1 cluster was specific to TNW1 within the analysed dataset. Species-level analysis identified 10 core, 20 variables, and 5 single-genome clusters in *F. avenaceum*, compared with 11 core, 9 variable, and 2 single-genome clusters in *F. graminearum* (Fig. 6). Detailed comparisons are provided in supplementary File S6.

**Fig. 5.**
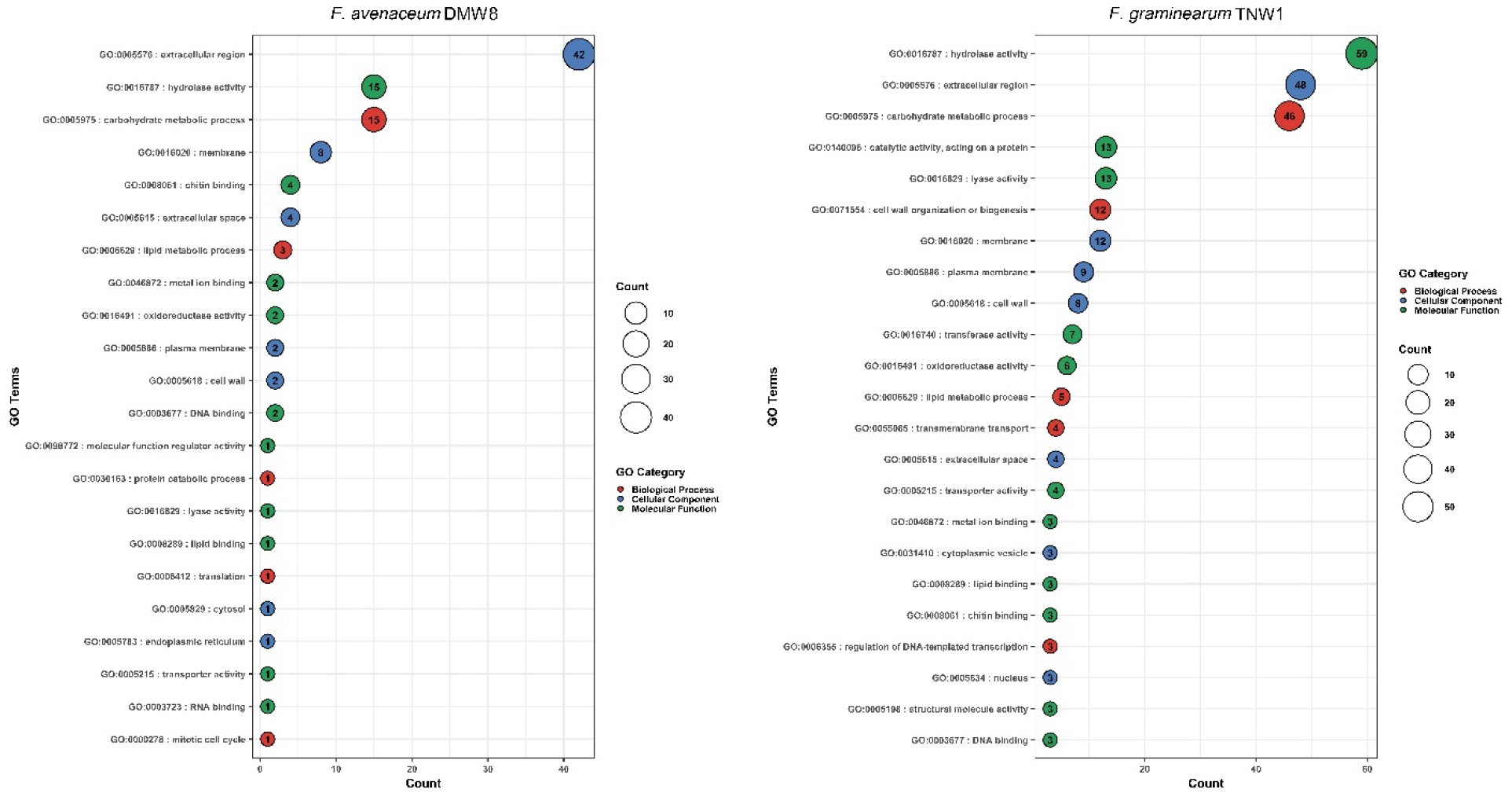
Bubble plot showing the top 10 Gene Ontology (GO) functional categories identified among predicted effector proteins across the analysed genomes. Bubble size represents the number of effector proteins assigned to each GO category, and bubble colour indicates the corresponding GO category or functional class.

**Fig. 6.**
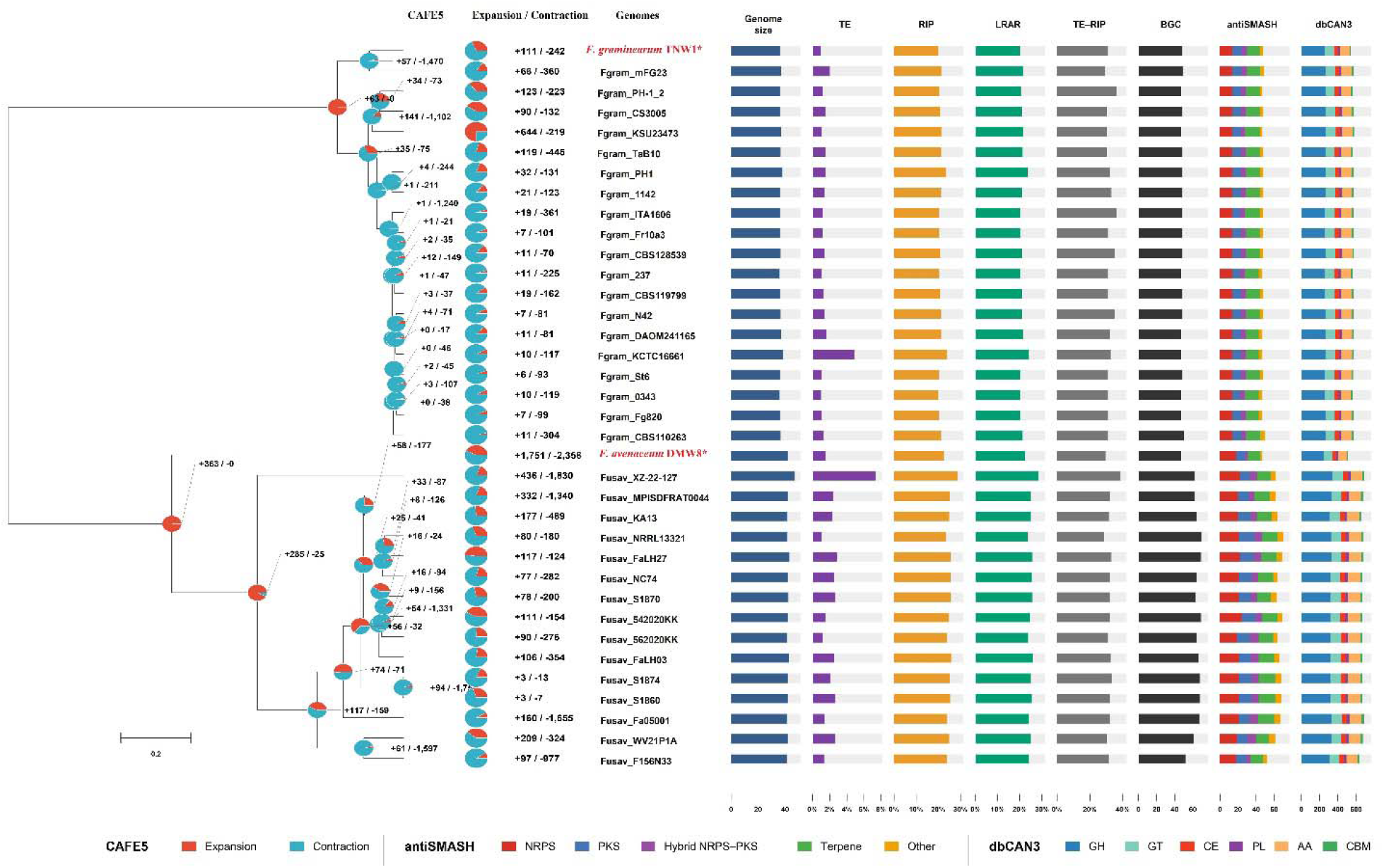
Integrated phylogenomic and genomic characterization of 36 Fusarium genomes. CAFE5-predicted gene-family expansions (red) and contractions (blue) are shown as pie charts. Genome-level features include genome size, TEs, RIPs, LRARs, TE–RIP overlap, and total BGCs. Stacked bars show antiSMASH BGC classes (NRPS, PKS, hybrid NRPS–PKS, terpene, and other) and dbCAN3 CAZyme classes (GH, GT, CE, PL, AA, and CBM). Asterisks indicate F. avenaceum DMW8 and F. graminearum TNW1.

### 4.7 Reduced CAZyme repertoires distinguish TNW1 and DMW8 from their respective reference genomes

The newly sequenced isolates exhibited distinct CAZyme repertoires relative to their respective species groups. *F. graminearum* TNW1 contained 538 CAZy annotations, representing the lowest total CAZyme count among the analysed *F. graminearum* genomes (560–579), with reduced GH (255), CE (42), AA (107), and particularly CBM (8) complements relative to the reference genomes. Notably, the CBM repertoire of TNW1 was substantially lower than that of the analysed *F. graminearum* genomes, which contained 16–21 CBMs. *F. avenaceum* (DMW8) displayed a more pronounced reduction, with 511 total CAZy annotations compared with 640–694 in the reference genomes. *F. avenaceum* (DMW8) had the lowest representation across all six major CAZy classes, including GH (248), GT (95), CE (39), PL (24), AA (98), and CBM (7), compared with ranges of 305–342, 107–114, 52–58, 26–29, 124–138, and 21–26, respectively (supplementary file S7), among the analysed *F. avenaceum* genomes (Fig. 6). Thus, while *F. graminearum* TNW1 was particularly distinguished by its reduced CBM complement, DMW8 exhibited a broad reduction across the CAZy repertoire. (Table 3)

**Table 3.** Comparative distribution of carbohydrate-active enzymes (CAZymes) in analysed genomes, showing total CAZyme content and classification into glycoside hydrolases (GH), glycosyltransferases (GT), auxiliary activities (AA), carbohydrate esterases (CE), polysaccharide lyases (PL), and carbohydrate-binding modules (CBM), along with the number of predicted secretory CAZymes.

| **Species** | **Genome accession number** | **AA** | **CBM** | **CE** | **GH** | **GT** | **PL** |
|---|---|---|---|---|---|---|---|
| *F. graminearum* TNW1 | GCA_054643435.1 | 107 | 8 | 42 | 255 | 102 | 24 |
| *F. avenaceum* DMW8 | GCA_054643395.2 | 98 | 7 | 39 | 248 | 95 | 24 |
| *F. graminearum*_0343 | GCA_018345745.1 | 113 | 16 | 47 | 257 | 104 | 23 |
| *F. graminearum*_1142 | GCA_018346805.1 | 114 | 18 | 48 | 265 | 105 | 24 |
| *F. graminearum*_237 | GCA_018346265.1 | 114 | 18 | 45 | 259 | 104 | 23 |
| *F. graminearum*_CBS110263 | GCA_018345365.1 | 114 | 17 | 48 | 262 | 104 | 24 |
| *F. graminearum*_CBS119799 | GCA_018345325.1 | 115 | 17 | 47 | 261 | 105 | 24 |
| *F. graminearum*_CBS128539 | GCA_018346575.1 | 116 | 18 | 48 | 264 | 106 | 24 |
| *F. graminearum* DAOM241165 | GCA_000966645.2 | 115 | 18 | 48 | 265 | 105 | 24 |
| *F. graminearum*_Fg820 | GCA_018346565.1 | 115 | 18 | 48 | 262 | 105 | 24 |
| *F. graminearum*_Fr10a3 | GCA_018346705.1 | 115 | 18 | 48 | 261 | 106 | 24 |
| *F. graminearum*_ITA1606 | GCA_018345645.1 | 115 | 18 | 48 | 260 | 106 | 23 |
| *F. graminearum*_N42 | GCA_018346015.1 | 115 | 18 | 48 | 264 | 105 | 24 |
| *F. graminearum*_PH1 | GCA_020991245.1 | 115 | 19 | 46 | 264 | 104 | 24 |
| *F. graminearum*_St6 | GCA_018345585.1 | 115 | 18 | 46 | 261 | 103 | 24 |
| *F. graminearum*_mFG23 | GCA_051903665.1 | 117 | 17 | 47 | 262 | 106 | 24 |
| *F. avenaceum*_542020KK | GCA_019055345.1 | 133 | 24 | 54 | 319 | 109 | 29 |
| *F. avenaceum*_562020KK | GCA_019055315.1 | 133 | 24 | 53 | 323 | 110 | 28 |
| *F. avenaceum*_F156N33 | GCA_018282135.1 | 124 | 21 | 55 | 305 | 108 | 27 |
| *F. avenaceum*_Fa05001 | GCA_000769215.1 | 135 | 26 | 55 | 332 | 111 | 28 |
| *F. avenaceum*_FaLH03 | GCA_000769305.1 | 131 | 26 | 55 | 321 | 110 | 29 |
| *F. avenaceum*_FaLH27 | GCA_000769295.1 | 135 | 25 | 55 | 329 | 109 | 29 |
| *F. avenaceum*_KA13 | GCA_012959155.1 | 129 | 22 | 54 | 313 | 110 | 27 |
| *F. avenaceum* MPISDFRAT0044 | GCA_020744115.1 | 129 | 25 | 56 | 331 | 110 | 27 |
| *F. avenaceum*_NC74 | GCA_044646885.1 | 134 | 23 | 55 | 319 | 107 | 29 |
| *F. avenaceum*_NRRL13321 | GCA_013753855.1 | 138 | 24 | 54 | 326 | 110 | 29 |
| *F. avenaceum*_S1860 | GCA_019055295.1 | 135 | 24 | 52 | 321 | 111 | 27 |
| *F. avenaceum*_S1870 | GCA_019055285.1 | 133 | 23 | 53 | 324 | 107 | 29 |
| *F. avenaceum*_S1874 | GCA_019055275.1 | 135 | 24 | 52 | 321 | 111 | 27 |
| *F. avenaceum*_WV21P1A | GCA_025948275.1 | 132 | 25 | 55 | 328 | 107 | 27 |
| *F. avenaceum*_XZ-22-127 | GCA_057929635.1 | 131 | 23 | 58 | 342 | 114 | 26 |
| *F. graminearum*_CS3005 | GCA_000599445.1 | 113 | 20 | 48 | 266 | 105 | 24 |
| *F. graminearum*_KCTC16661 | GCA_055476185.1 | 116 | 18 | 48 | 264 | 105 | 24 |
| *F. graminearum*_KSU23473 | GCA_037255895.1 | 109 | 21 | 47 | 270 | 108 | 24 |
| *F. graminearum*_PH-1_2 | GCF_000240135.3 | 111 | 21 | 46 | 262 | 102 | 24 |
| *F. graminearum*_TaB10 | GCA_012959185.1 | 112 | 17 | 48 | 262 | 101 | 24 |

### 4.8 Low transposable-element content and RIP-associated coverage characterise both Indian isolates

The newly sequenced Indian *F. graminearum* isolate TNW1 showed a relatively low TE content compared with the other *F. graminearum* isolates analysed, with TEs occupying 0.96% of the genome. This was comparable to Fgram_0343 (1.00%) and Fgram_237 (1.10%). Among the analysed *F. graminearum* genomes, TNW1 therefore exhibited one of the lower levels of TE accumulation (Fig. S3).

Similarly, the newly sequenced Indian *F. avenaceum* isolate DMW8 contained 1.55% TE content, placing it toward the lower end of the *F. avenaceum* genomes analysed. TE content among the *F. avenaceum* isolates ranged from 1.07% in Fusav_NRRL13321 to 7.39% in Fusav_XZ-22-127, with DMW8 falling within the lower portion of this distribution (Table 4). Overall, TNW1 and DMW8 exhibited relatively low TE accumulation within their respective species groups (Fig. 6).

**Table 4.** Comparison of genome size, RIP signatures, LRARs, and TE content across 36 analysed genomes. Values include RIP extent, LRAR count and length, and TE copy number, total length, density, and mean size

| **Genome** | **Genome size in Mb** | **Scaffold/ contig count** | **TE interval count** | **TE total length Mb** | **TE percent genome** | **RIP window count** | **RIP total length mb** | **RIP % genome** | **LRAR count** | **LRAR total length bp** | **LRAR % genome** | **RIP TE overlap records** | **LRAR with TE overlap %** |
|---|---|---|---|---|---|---|---|---|---|---|---|---|---|
| *F. graminearum* TNW1 | 36.29 | 21 | 7001 | 0.35 | 0.96 | 4871 | 7.26 | 20.01 | 4871 | 7.26 | 20.01 | 1499 | 30.77 |
| *F. avenaceum* DMW8 | 42.00 | 35 | 8285 | 0.65 | 1.55 | 6082 | 9.40 | 22.38 | 6082 | 9.40 | 22.38 | 1792 | 29.46 |
| *F. graminearum*_0343 | 36.07 | 17 | 6969 | 0.36 | 1.00 | 4905 | 7.23 | 20.03 | 4905 | 7.23 | 20.03 | 1515 | 30.89 |
| *F. graminearum*_1142 | 36.66 | 17 | 7467 | 0.51 | 1.39 | 4893 | 7.72 | 21.06 | 4893 | 7.72 | 21.06 | 1612 | 32.95 |
| *F. graminearum*_237 | 36.00 | 20 | 7013 | 0.40 | 1.10 | 4856 | 7.30 | 20.28 | 4856 | 7.30 | 20.28 | 1507 | 31.03 |
| *F. graminearum*_CBS110263 | 36.77 | 34 | 7296 | 0.47 | 1.27 | 4980 | 7.79 | 21.19 | 4980 | 7.79 | 21.19 | 1541 | 30.94 |
| *F. graminearum*_CBS119799 | 36.66 | 46 | 7205 | 0.46 | 1.25 | 4961 | 7.66 | 20.89 | 4961 | 7.66 | 20.89 | 1531 | 30.86 |
| *F. graminearum*_CBS128539 | 36.71 | 35 | 7673 | 0.51 | 1.39 | 4992 | 7.66 | 20.86 | 4992 | 7.66 | 20.86 | 1744 | 34.94 |
| *F. graminearum*_DAOM241165 | 37.02 | 22 | 7421 | 0.60 | 1.62 | 4971 | 7.88 | 21.29 | 4971 | 7.88 | 21.29 | 1592 | 32.03 |
| *F. graminearum*_Fg820 | 36.39 | 20 | 6971 | 0.38 | 1.04 | 4933 | 7.35 | 20.19 | 4933 | 7.35 | 20.19 | 1534 | 31.10 |
| *F. graminearum*_Fr10a3 | 36.42 | 32 | 7076 | 0.43 | 1.17 | 4912 | 7.42 | 20.38 | 4912 | 7.42 | 20.38 | 1530 | 31.15 |
| *F. graminearum*_ITA1606 | 36.26 | 32 | 7748 | 0.44 | 1.22 | 4878 | 7.31 | 20.16 | 4878 | 7.31 | 20.16 | 1751 | 35.90 |
| *F. graminearum*_N42 | 36.78 | 9 | 7840 | 0.51 | 1.38 | 4936 | 7.76 | 21.10 | 4936 | 7.76 | 21.10 | 1723 | 34.91 |
| *F. graminearum*_PH1 | 38.04 | 5 | 7363 | 0.59 | 1.55 | 5090 | 8.92 | 23.45 | 5090 | 8.92 | 23.45 | 1637 | 32.16 |
| *F. graminearum*_St6 | 36.19 | 24 | 7012 | 0.38 | 1.05 | 4921 | 7.31 | 20.20 | 4921 | 7.31 | 20.20 | 1509 | 30.66 |
| *F. graminearum*_mFG23 | 37.05 | 5 | 7256 | 0.74 | 2.01 | 4966 | 7.95 | 21.46 | 4966 | 7.95 | 21.46 | 1446 | 29.12 |
| *F. avenaceum*_542020KK | 42.35 | 177 | 8381 | 0.62 | 1.47 | 6303 | 10.42 | 24.61 | 6303 | 10.42 | 24.61 | 2033 | 32.25 |
| *F. avenaceum*_562020KK | 41.51 | 129 | 7786 | 0.50 | 1.20 | 6192 | 9.88 | 23.80 | 6192 | 9.88 | 23.80 | 1921 | 31.02 |
| *F. avenaceum*_F156N33 | 41.19 | 214 | 8040 | 0.55 | 1.33 | 6166 | 9.84 | 23.88 | 6166 | 9.84 | 23.88 | 1947 | 31.58 |
| *F. avenaceum*_Fa05001 | 41.59 | 83 | 8034 | 0.60 | 1.43 | 6201 | 9.97 | 23.98 | 6201 | 9.97 | 23.98 | 1971 | 31.79 |
| *F. avenaceum*_FaLH03 | 42.71 | 105 | 8759 | 1.08 | 2.52 | 6245 | 11.03 | 25.82 | 6245 | 11.03 | 25.82 | 2049 | 32.81 |
| *F. avenaceum*_FaLH27 | 43.17 | 78 | 9128 | 1.24 | 2.88 | 6290 | 11.06 | 25.61 | 6290 | 11.06 | 25.61 | 2070 | 32.91 |
| *F. avenaceum*_KA13 | 41.70 | 34 | 8456 | 0.97 | 2.34 | 6054 | 10.39 | 24.91 | 6054 | 10.39 | 24.91 | 1923 | 31.76 |
| *F. avenaceum*_MPISDFRAT0044 | 42.23 | 13 | 8477 | 1.03 | 2.43 | 6177 | 10.55 | 24.99 | 6177 | 10.55 | 24.99 | 1974 | 31.96 |
| *F. avenaceum*_NC74 | 42.28 | 68 | 8511 | 1.04 | 2.47 | 6191 | 10.75 | 25.43 | 6191 | 10.75 | 25.43 | 1975 | 31.90 |
| *F. avenaceum*_NRRL13321 | 41.72 | 778 | 7598 | 0.45 | 1.07 | 6221 | 9.79 | 23.47 | 6221 | 9.79 | 23.47 | 1762 | 28.32 |
| *F. avenaceum*_S1860 | 42.33 | 20 | 8866 | 1.13 | 2.66 | 6099 | 10.73 | 25.36 | 6099 | 10.73 | 25.36 | 1968 | 32.27 |
| *F. avenaceum*_S1870 | 42.46 | 29 | 8675 | 1.13 | 2.65 | 6123 | 10.85 | 25.56 | 6123 | 10.85 | 25.56 | 1958 | 31.98 |
| *F. avenaceum*_S1874 | 42.10 | 139 | 8632 | 0.89 | 2.11 | 6229 | 10.51 | 24.98 | 6229 | 10.51 | 24.98 | 2080 | 33.39 |
| *F. avenaceum*_WV21P1A | 42.04 | 9 | 8035 | 1.11 | 2.65 | 6129 | 10.47 | 24.90 | 6129 | 10.47 | 24.90 | 1872 | 30.54 |
| *F. avenaceum*_XZ-22-127 | 46.98 | 9 | 12039 | 3.47 | 7.39 | 7012 | 13.41 | 28.55 | 7012 | 13.41 | 28.55 | 2693 | 38.41 |
| Fusgr_CS3005 | 36.67 | 199 | 6914 | 0.54 | 1.47 | 5011 | 7.65 | 20.86 | 5011 | 7.65 | 20.86 | 1516 | 30.25 |
| Fusgr_KCTC16661 | 38.35 | 4 | 7191 | 1.87 | 4.87 | 5163 | 9.19 | 23.97 | 5163 | 9.19 | 23.97 | 1687 | 32.67 |
| Fusgr_KSU23473 | 36.88 | 5 | 7274 | 0.38 | 1.04 | 4949 | 7.92 | 21.48 | 4949 | 7.92 | 21.48 | 1507 | 30.45 |
| Fusgr_PH-1_2 | 36.46 | 19 | 7645 | 0.44 | 1.21 | 4949 | 7.48 | 20.51 | 4949 | 7.48 | 20.51 | 1780 | 35.97 |
| Fusgr_TaB10 | 36.76 | 54 | 7156 | 0.55 | 1.50 | 4954 | 7.80 | 21.22 | 4954 | 7.80 | 21.22 | 1507 | 30.42 |

### Repeat-induced point mutation (RIP) patterns

RIP analysis revealed variation in genome-wide RIP-associated coverage across the Fusarium genomes analysed. The Indian isolates showed comparatively lower RIP-associated coverage within their respective species groups, with 20.01% in TNW1 and 22.38% in DMW8. TNW1 contained 4,871 RIP-associated windows, corresponding to 7.26 Mb of the genome, whereas DMW8 contained 6,082 RIP-associated windows covering 9.40 Mb.

Among the *F. graminearum* isolates, RIP-associated coverage ranged from 20.01% in TNW1 to 23.45% in Fgram_PH1, with TNW1 showing the lowest value. Similarly, among the *F. avenaceum* isolates, RIP-associated coverage ranged from 22.38% in DMW8 to 28.55% in Fusav_XZ-22-127, with DMW8 showing the lowest value (Table 4). Overall, both Indian isolates exhibited RIP-associated coverage toward the lower end of the distributions observed within their respective species groups (Fig. 6).

### TE insertion dynamics in Indian Fusarium isolates

TE insertion analysis using the McClintock pipeline identified candidate TE insertion sites in both Indian isolates. TEFLoN identified 512 reference-associated insertions in TNW1 and 593 candidate insertions in DMW8, comprising 591 reference-associated and two non-reference insertions. These results describe the insertion patterns detected by the selected method and do not independently demonstrate ongoing transposition.

Across detection methods, TEMP identified 723 insertions in TNW1 and 788 in DMW8, while ngs_te_mapper2 detected 173 and 159 insertions, respectively. Sequencing coverage was 22.3× for TNW1 and 28.1× for DMW8. Across detection methods, candidate insertion counts varied among isolates, with DMW8 showing higher counts than TNW1 in some analyses. These differences do not, by themselves, establish differences in current TE activity. (Table 5).

**Table 5.** Summary of transposable element (TE) insertions detected in DMW8 and TNW1 genomes using the McClintock pipeline.

| **Category** | **Metric / Method** | ***F. avenaceum* DMW8** | ***F. graminearum* TNW1** |
|---|---|---|---|
| Read & Coverage Statistics | Read Length | 151 | 151 |
|  | Reads (R1) | 18458384 | 13751992 |
|  | Reads (R2) | 18459160 | 13752190 |
|  | Insert Size | 121 | 121 |
|  | Coverage (x) | 28.09 | 22.3 |
| Transposable Element Insertion Calls (McClintock) | ngs_te_mapper (ALL) | 2 | 7 |
|  | ngs_te_mapper (REF) | 2 | 7 |
|  | ngs_te_mapper (NONREF) | 0 | 0 |
|  | ngs_te_mapper2 (ALL) | 159 | 173 |
|  | ngs_te_mapper2 (REF) | 159 | 172 |
|  | ngs_te_mapper2 (NONREF) | 0 | 1 |
|  | relocate (ALL) | 93 | 102 |
|  | relocate (REF) | 93 | 102 |
|  | relocate (NONREF) | 0 | 0 |
|  | relocate2 (ALL) | 126 | 99 |
|  | relocate2 (REF) | 126 | 97 |
|  | relocate2 (NONREF) | 0 | 2 |
|  | temp (ALL) | 788 | 723 |
|  | temp (REF) | 788 | 723 |
|  | temp (NONREF) | 0 | 0 |
|  | temp2 (ALL) | 788 | 723 |
|  | temp2 (REF) | 788 | 723 |
|  | temp2 (NONREF) | 0 | 0 |
|  | retroseq (ALL) | 0 | 0 |
|  | retroseq (REF) | 0 | 0 |
|  | retroseq (NONREF) | 0 | 0 |
|  | popoolationte (ALL) | 1 | 1 |
|  | popoolationte (REF) | 1 | 0 |
|  | popoolationte (NONREF) | 0 | 1 |
|  | popoolationte2 (ALL) | 62 | 40 |
|  | popoolationte2 (REF) | 61 | 39 |
|  | popoolationte2 (NONREF) | 1 | 1 |
|  | te-locate (ALL) | 0 | 0 |
|  | te-locate (REF) | 0 | 0 |
|  | te-locate (NONREF) | 0 | 0 |
|  | teflon (ALL) | 593 | 512 |
|  | teflon (REF) | 591 | 512 |
|  | teflon (NONREF) | 2 | 0 |
|  | tebreak (ALL) | 1 | 0 |
|  | tebreak (REF) | 0 | 0 |
|  | tebreak (NONREF) | 1 | 0 |

### 4.9 Contrasting orthogroup composition and gene-family turnover in TNW1 and DMW8

A total of 11,427 gene clusters and 87 singletons were identified in *F. graminearum* TNW1, whereas 10,708 gene clusters and 693 singletons were identified in *F. avenaceum* DMW8. Among the other genomes, 12,177 gene clusters and 475 singletons were identified in *F. avenaceum* (FaLH03), 11,973 gene clusters and 571 singletons in *F. avenaceum* (WV21P1A), 11,938 gene clusters and 1,002 singletons in *F. graminearum*, 11,316 gene clusters and 1,199 singletons in *F. poae*, 11,271 gene clusters and 59 singletons in *F. graminearum* (mFG23), 11,140 gene clusters and 125 singletons in *F. asiaticum* (KCTC16664), 10,968 gene clusters and 53 singletons in *F. boothii* (CBS31673), and 10,878 gene clusters and 331 singletons in *F. culmorum*. (Table 6). All ten genomes together formed 14,975 gene clusters, 540 overlapping clusters, 6,146 single-copy gene clusters, and 4,595 singletons (Table 6). The total number of proteins across all genomes was 123,738, with singletons accounting for 3.71% of the total protein repertoire (Fig. 7a).

**Fig. 7.**
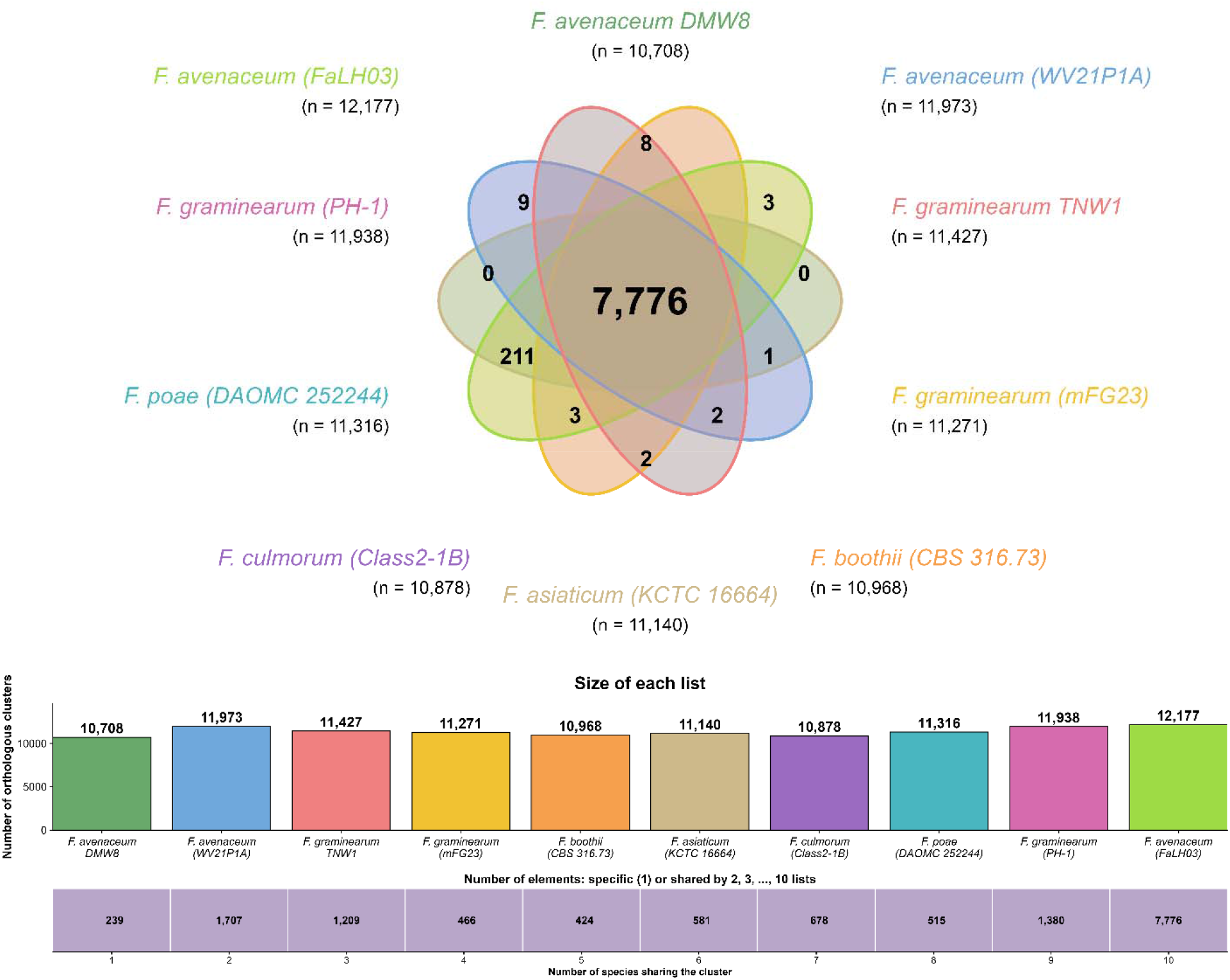
**(a)** Comparative analysis of shared and unique orthologous gene clusters among ten Fusarium genomes, including the newly assembled genomes Fusarium graminearum TNW1 and Fusarium avenaceum DMW8, generated using OrthoVenn3.

**Table 6.** Orthologous clustering analysis of predicted proteomes from selected *Fusarium* species, including cluster distribution, singleton representation, and overall clustering metrics.

| **6a. Species Statistics** |  |  |  |
| **Species** | **Proteins** | **Clusters** | **Singletons** |
|---|---|---|---|
| *F. poae* | 13,851 | 11,316 | 1,199 |
| *F. graminearum* | 13,312 | 11,938 | 1,002 |
| *F. avenaceum* DMW8 | 13,028 | 10,708 | 693 |
| *F. avenaceum* FaLH03 | 13,009 | 12,177 | 475 |
| *F. avenaceum* WV21P1A | 12,876 | 11,973 | 571 |
| *F. graminearum* TNW1 | 11,750 | 11,427 | 87 |
| *F. graminearum* mFG23 | 11,550 | 11,271 | 59 |
| *F. asiaticum* KCTC16664 | 11,504 | 11,140 | 125 |
| *F. culmorum* | 11,444 | 10,878 | 331 |
| *F. boothii* CBS31673 | 11,414 | 10,968 | 53 |
| **6b. Distribution of unique clusters associated with the virulence and pathogenicity shared by 10 Fusarium species analysed.** |  |  |  |
| **Pathway** | **Cluster Count** | **Percentage (%)** |  |
| MAPK_HOG_CWI | 33 | **8.99%** |  |
| cAMP_Gprotein | 6 | **1.63%** |  |
| TOR_pathway | 72 | **19.62%** |  |
| Calcium_Calcineurin | 1 | **0.27%** |  |
| Transcription_factors | 101 | **27.52%** |  |
| Secondary_metabolism | 43 | **11.72%** |  |
| CWDE | 62 | **16.89%** |  |
| Virulence_core | 49 | **13.35%** |  |
| **Total** | **367** | **99.99% ≈ 100%** |  |
| **6c. Summary of orthologous clusters shared by 10 Fusarium species analysed.** |  |  |  |
| **Metric** | **Value** |  |  |
| Overlaps | 540 |  |  |
| All clusters | 14,975 |  |  |
| Single-copy clusters | 6,146 |  |  |
| All proteins | 123,738 |  |  |
| All singletons | 4,595 |  |  |
| Percentage of singletons | 3.71% |  |  |

Among these, 7,776 orthologous gene clusters were shared across all ten genomes. A total of 239 clusters were uniquely associated with a single genome, of which 211 were specific to *F. poae*, followed by nine in *F. avenaceum* (FaLH03), eight in DMW8, three each in *F. culmorum* and *F. avenaceum* (WV21P1A), two each in *F. boothii* (CBS31673) and *F. asiaticum* (KCTC16664), and one in *F. graminearum* (mFG23). No unique clusters were detected in TNW1 or the *F. graminearum* reference genome. Pairwise sharing revealed that *F. avenaceum* (FaLH03) shared 846 clusters with *F. avenaceum* (WV21P1A), followed by *F. culmorum* with *F. graminearum*, which shared 238 clusters (Fig. 7a, 7b). DMW8 shared 82 clusters with *F. avenaceum* (FaLH03) and 63 with *F. avenaceum* (WV21P1A), whereas *F. graminearum* shared 60 clusters with *F. poae*. Furthermore, *F. graminearum* (mFG23) shared 58 clusters with TNW1, while *F. culmorum* shared 47 clusters with *F. poae,* and *F. poae* shared 40 clusters with *F. avenaceum* (FaLH03) (S1).

**Fig. 7.**
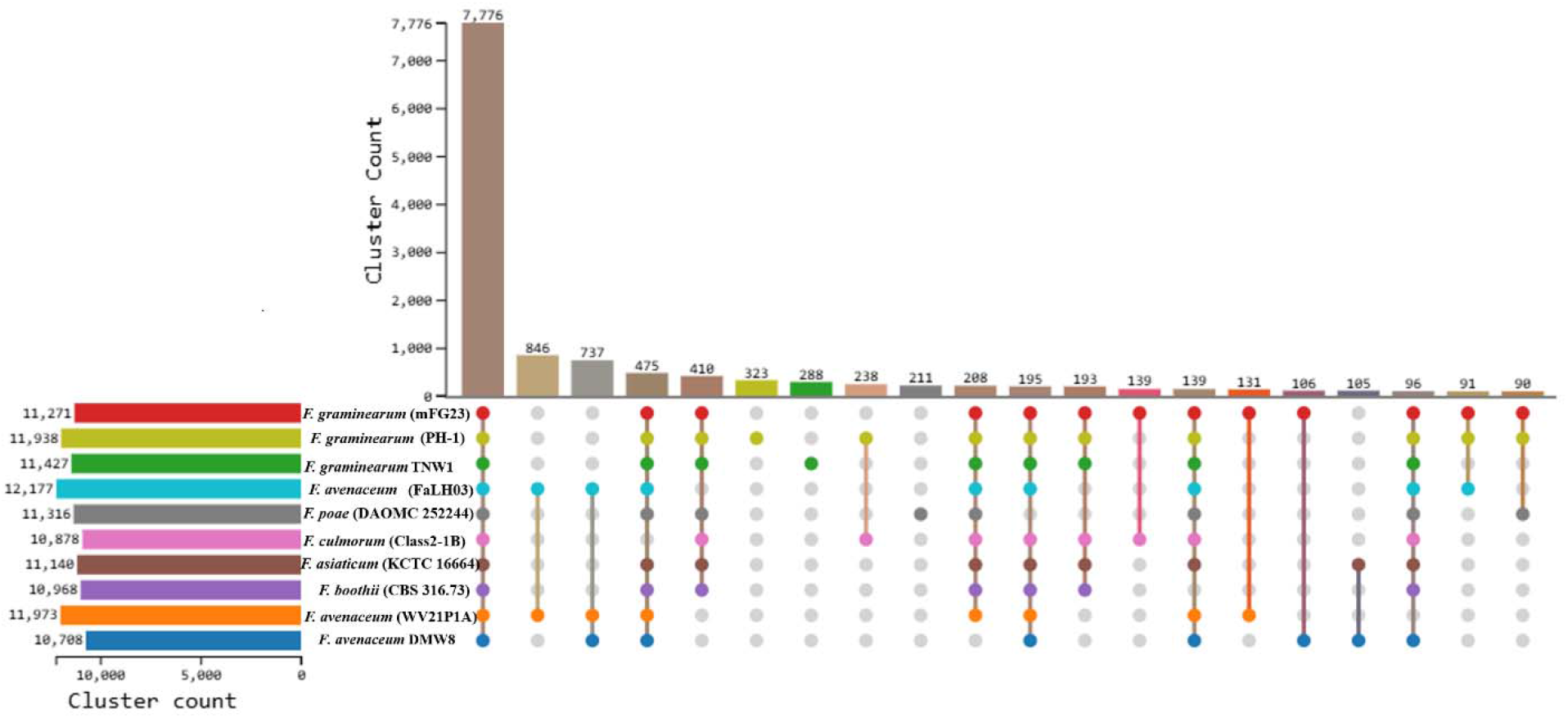
**(b)** UpSet plot illustrating the intersections of orthologous gene clusters among the ten Fusarium genomes analysed.

Gene-family expansion and contraction analysis using CAFE5 revealed contrasting patterns among the *Fusarium* genomes in the 10-genome dataset (Fig. 8). DMW8 showed 1,767 expanded and 1,273 contracted gene families, whereas TNW1 exhibited 98 expansions and 258 contractions. Among the reference genomes, *F. culmorum* showed the highest number of contracted gene families (1,314), while the other genomes displayed varying expansion–contraction patterns. The corresponding results for the 36-genome dataset are presented separately (Fig. 8).

**Fig. 8.**
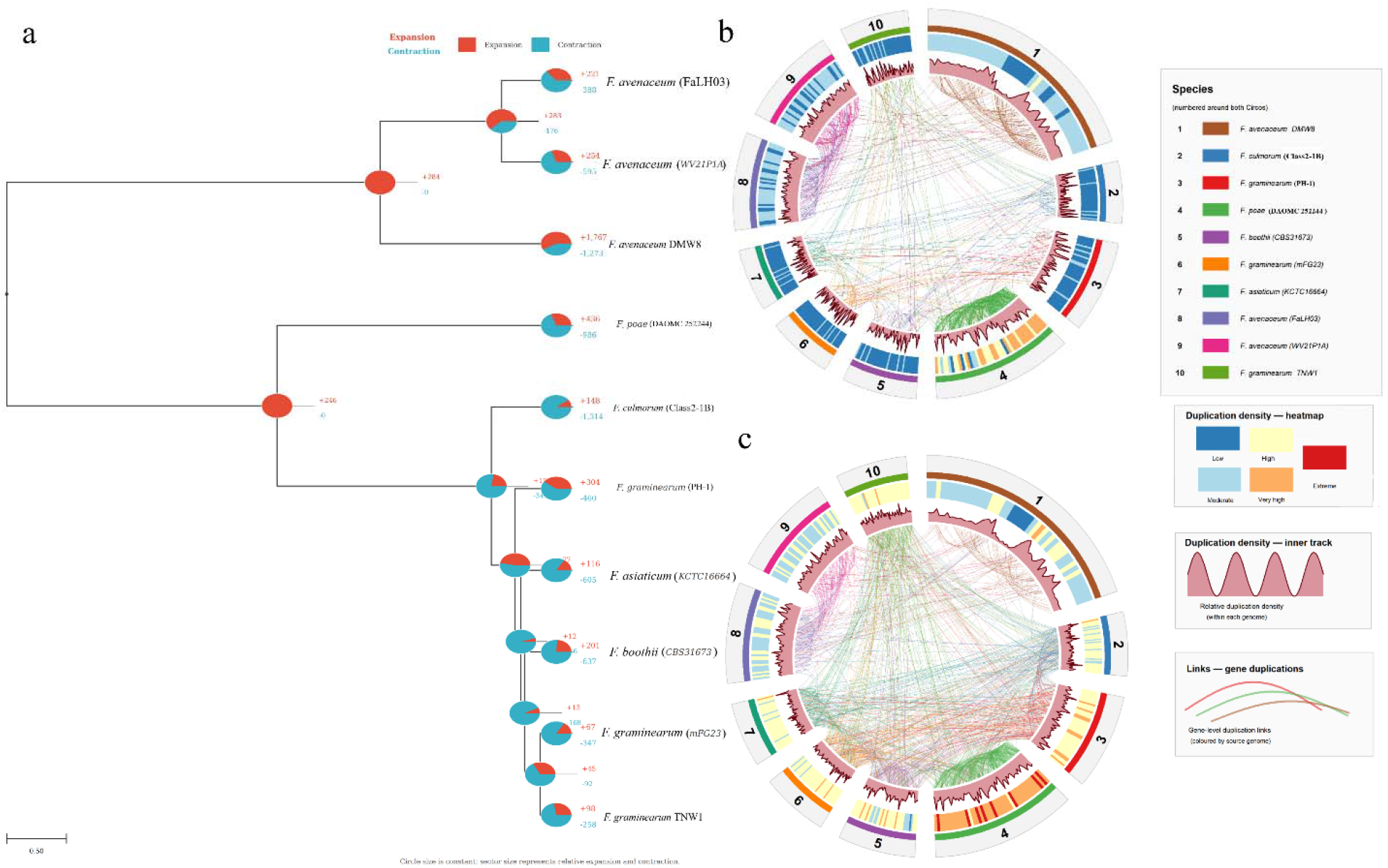
Comparative orthology and gene duplication analysis of ten Fusarium genomes based on OrthoFinder results. (a) Phylogenetic relationships and expansion and contraction of orthologous gene families among the analysed Fusarium genomes. Pie charts at the nodes represent expanded (red) and contracted (blue) gene families, with the corresponding numbers of expanded and contracted families indicated beside each node. (b) Circos plot showing high-confidence gene duplication events (support ≥0.98) across the analysed genomes. The numbered outer sectors correspond to the genomes listed in the species legend. The heatmap track represents duplication density, ranging from low to extreme, while the inner histogram track shows relative duplication density within each genome. Coloured links represent gene-level duplication relationships, with link colours indicating the source genome. (c) Circos plot showing all detected gene duplication events across the analysed genomes using the same genome sectors and duplication-density tracks as in panel without applying the high-confidence support threshold.

A total of 367 pathway-associated orthologous clusters were identified across all genomes. These were mainly associated with transcription factors (27.52%), followed by TOR signalling (19.62%), cell wall-degrading enzymes (16.89%), and virulence core pathways (13.35%). Smaller contributions were observed from secondary metabolism (11.72%), MAPK–HOG–CWI signalling (8.99%), cAMP/G-protein signalling (1.63%), and calcium–calcineurin signalling (0.27%) (Table 6).

Analysis of the 10-genome dataset identified 13,909 orthogroups, including 7,859 shared across all genomes, with 97.8% of predicted proteins assigned to orthogroups (Fig. 7b). A total of 5,489 single-copy orthogroups and 155 species-specific orthogroups were detected. OrthoFinder identified 6,209 duplication events, including 4,876 high-confidence events (≥0.98). DMW8 showed the highest proportion of genes in duplicated orthogroups (41.71%), whereas TNW1 showed 33.23%, indicating differences in gene copy-number profiles among the analysed genomes (Fig. 7b).

In the 36-genome analysis, 16,504 orthogroups containing 441,387 genes were identified, including 7,625 orthogroups shared across all genomes and 5,035 single-copy orthogroups. DMW8 shared 11,161–12,303 orthologues with other *F. avenaceum* genomes, while TNW1 shared 10,876–11,036 orthologues with other *F. graminearum* genomes. Overall orthogroup profiles showed strong conservation across both species groups, with greater similarity among genomes from the same species (Fig. S4). Pairwise Jaccard similarity based on orthogroup presence–absence profiles further supported this pattern, showing higher similarity within species groups than between *F. graminearum* and *F. avenaceum* genomes (Fig. S5).

DMW8 exhibited 1,664 terminal duplication events, compared with 53 in TNW1. In the 36-genome CAFE5 analysis, DMW8 showed 1,751 expanded and 2,356 contracted gene families, whereas TNW1 showed 111 expansions and 242 contractions. Despite this higher turnover, DMW8 contained only one genome-specific orthogroup (two genes), whereas none were detected in TNW1. Detailed orthologue, duplication, and gene-family expansion/contraction are shown in Fig. 8, and the results are provided in Supplementary File S8.

## 5. Discussion

Genome sequencing of plant pathogens has significantly advanced our understanding of pathogen biology, genome evolution, and host–pathogen interactions. Fusarium head blight (FHB) remains an important concern under changing agro-climatic conditions; however, genome-level information on Indian *Fusarium* isolates remains limited. Fungal genome evolution is shaped by interactions among mobile genetic elements, genome defence mechanisms, ecological pressures, and lifestyle transitions (Kelkar & Ochman, 2012; Stukenbrock, 2013; Gladyshev et al., 2017; Muszewska et al., 2019), highlighting the importance of region-specific genomic studies for understanding pathogen adaptation and evolutionary dynamics. In this study, we present high-quality genome assemblies of *F. graminearum* TNW1 and *F. avenaceum* DMW8 isolated from symptomatic wheat tissues in India. Hybrid sequencing using Illumina and Oxford Nanopore platforms generated highly contiguous assemblies, with genome sizes of 36.29 Mb for TNW1 and 42 Mb for DMW8, respectively, with high completeness as assessed by BUSCO. The genome sizes of both isolates fall within the ranges reported for their respective species. High sequencing depth, coverage, and completeness further supported the quality and robustness of the assemblies.

In the Indian context, genomic resources for *F. graminearum* and *F. avenaceum* are not yet available, highlighting a critical gap in region-specific pathogen genomics. The most recent wheat-derived genomes available for *F. graminearum* are from (DeGenring et al., 2026), infectious on *T. turgidum var. durum* from Ethiopia, whereas F. avenaceum lacks recent wheat-associated genome sequences, with available data limited to 2014 (Lysøe et al., 2014) using 454 pyrosequencing and Illumina short-read technologies, with limited functional and comparative analysis. Newer submissions originate from non-wheat hosts such as *Pinus taeda*. (accessed 10 July 2026). The genome assemblies generated in this study provide a genomic framework for Indian FHB-associated *F. graminearum* and *F. avenaceum*. The TNW1 genome (36.29 Mb) is within the reported genome-size range for *F. graminearum*, while the DMW8 assembly (42 Mb) is comparable to previously reported *F. avenaceum* genomes (approximately 41.6–43.1 Mb).

Comparative gene-family analysis revealed differences in gene-family turnover between TNW1 and DMW8 across the two datasets. TNW1 consistently exhibited more gene-family contractions than expansions, whereas DMW8 showed contrasting patterns, with more expansions in the broader 10-genome comparison but more contractions in the 36-genome comparison. These differences highlight the influence of taxonomic sampling and evolutionary context on inferred patterns of gene-family evolution. Gene duplication and loss are important processes contributing to changes in genome content and evolutionary diversification in plant-pathogenic fungi (Raffaele and Kamoun, 2012; Albalat and Cañestro, 2016). The observed contractions indicate reductions in gene-family copy numbers relative to the inferred ancestral state, although they do not necessarily represent complete gene-family loss or overall genome reduction.

Functional annotation identified genes associated with carbohydrate metabolism, transport and cellular signalling, including MAPK- and cAMP-associated pathways. These functions are relevant to fungal nutrient acquisition, environmental responses and interactions with plant hosts. In particular, carbohydrate-active enzymes contribute to the degradation of plant cell-wall components and the utilisation of plant-derived nutrients during fungal colonisation (Kubicek et al., 2014). However, the presence of genes assigned to these functional categories does not, by itself, demonstrate increased pathway activity or enhanced pathogenicity.

Phylogenomic analysis placed TNW1 within the F. graminearum clade and DMW8 within the F. avenaceum clade. Orthogroup profiles showed greater similarity among genomes belonging to the same species than between the two species groups. Shared functional annotations are consistent with conserved biological functions. This conservation is further supported by the predominance of shared virulence-associated pathways, particularly cAMP and MAPK signalling, which regulate key processes such as host infection, stress response, and toxin production.

Comparative KEGG analysis identified differences in the representation of signalling, secretion and host-interaction-related annotations in TNW1 and DMW8. These differences suggest candidate functional features for further investigation. In contrast, DMW8 exhibited enrichment in sesquiterpenoid and triterpenoid biosynthesis pathways, reflecting metabolic specialisation. The newly sequenced Indian isolate of *F. graminearum* harbours pathways for key virulence-associated mycotoxins, including trichothecenes (DON), zearalenone, fumonisins, and PKS14-linked orsellinic acid, indicating infection-related secondary metabolism and potential for food contamination. DON promotes host colonisation by suppressing plant defences, while in planta expression of PKS14 suggests orsellinic acid biosynthesis is infection-specific despite being undetectable in vitro (Jørgensen et al., 2014; Eskola et al., 2020; Armer et al., 2024). In contrast, Fusarium avenaceum produces metabolites such as beauvericin and enniatins, which are mainly associated with wheat grain contamination and toxicity (Logrieco et al., 2002). Overall, *F. graminearum* shows higher virulence, whereas *F. avenaceum* contributes primarily to contamination and moderate pathogenicity.

Secretome prediction identified candidate secreted proteins and effectors in both isolates, providing targets for future functional studies. The CAZyme analysis, however, showed differences between the isolates and their reference groups, particularly the reduced repertoire in DMW8. Together, the secretome, CAZyme and biosynthetic-cluster results indicate both shared predicted functions and differences in genomic potential; their consequences for infection and disease severity remain to be experimentally determined. Collectively, these findings suggest that pathogenicity in *Fusarium* is driven by conserved virulence mechanisms complemented by selective metabolic adaptations, enabling different species to cause Fusarium Head Blight in wheat (Ma et al., 2013; Kazan et al., 2012). Pathogenicity assays revealed disease-associated symptoms on wheat spikes and seeds, while histopathological observations provided additional evidence of fungal development, complementing the genomic characterisation of TNW1 and DMW8.

The Indian isolates of *F. graminearum* (TNW1) and *F. avenaceum* (DMW 8) possess highly compact repeat landscapes, characterised by markedly reduced transposable element (TE) content and density compared with other Fusarium species, such as *F. poae* and *F. culmorum*. Transposable elements are widely recognised as major drivers of genome evolution, structural variation, and ecological adaptation in fungi through their ability to mediate insertions, recombination, and genome expansion (Belyayev, 2014; Hess et al., 2014). However, fungal genomes display striking variability in TE content, ranging from as little as 0.02% to nearly 30% of the genome (Castanera et al, 2016), indicating that TE accumulation is not uniform even among closely related taxa.

In contrast to TE-expanded genomes such as *Cenococcum geophilum*, where transposable elements can constitute more than 70% of the genome (Floudas et al, 2012), and RIP-dominated systems such as *Neurospora crassa*, where extensive repeat-induced point mutation (RIP) generates large numbers of highly mutated TE remnants (Galagan & Selker, 2004; Clutterbuck, 2011), TNW1 and DMW8 display both low TE abundance and minimal RIP signatures. This combination reflects a TE-constrained genome architecture, indicative of limited historical TE expansion and reduced genome defence-driven mutational pressure (Peter et al., 2016; Castanera et al., 2016). Consistent with this, RIP is a fungal genome defence mechanism that introduces C \ T mutations in duplicated sequences (Galagan & Selker, 2004; Clutterbuck, 2011). RIP-associated coverage was comparatively low in both Indian isolates compared with the other genomes analysed, with 20.01% of the TNW1 genome and 22.38% of the DMW8 genome within RIP-associated regions. Although the efficiency of RIP varies widely among fungal species, ranging from extensive TE inactivation in *Neurospora crassa* to weaker or incomplete activity in other fungi (Santana & Queiroz, 2015), the comparatively low RIP-associated coverage observed in TNW1 and DMW8 may be related to the relatively low abundance of repetitive elements in these genomes. However, RIP patterns can also be influenced by genomic context and the availability of duplicated sequences.

Recent studies further indicate that TE dynamics are strongly influenced by genomic context. In *Zymoseptoria tritici*, TE bursts have been associated with proximity to genes and GC-rich genomic regions, where elements exhibit reduced RIP signatures and can escape genome defence mechanisms (Oggenfuss & Croll, 2023). This highlights that TE activity is not solely governed by genome-wide defence systems but also by local genomic environments. TE distribution can vary with local genomic context. The present analyses, however, do not establish whether genomic environments promote or suppress TE activity in TNW1 or DMW8

In fungal genomes, TE insertions can affect nearby genes, depending on their location and functional consequences. Although TEs have been associated with genome plasticity in several plant pathogens (e,g *Cladosporium fulvum*), they have been shown to mediate the loss of key avirulence genes, facilitating evasion of host immunity. These findings underscore the role of TEs as drivers of genome plasticity and adaptive evolution in host–pathogen interactions.

Despite this overall genomic stability, TE insertion analysis produced different numbers of candidate insertions. TEFLoN identified 512 reference-associated insertions in TNW1 and 593 insertions in DMW8, including 591 reference-associated and two non-reference insertions. Similar patterns have been reported in fungi, where genome defence mechanisms restrict large-scale TE expansion while allowing low-frequency polymorphic insertions to persist (Castanera et al., 2016). Divergence profiles further supported a TE-quiescent state, with most elements appearing highly degenerated and lacking evidence of recent amplification. In contrast, TE-rich fungal genomes often exhibit multiple waves of transposon expansion and recent bursts of activity (Castanera et al., 2016). In many plant-pathogenic fungi, TE-rich and RIP-affected regions contribute to a “two-speed genome” organisation, in which rapidly evolving compartments harbour effector genes and facilitate host adaptation (Dong et al., 2015; Rouxel et al., 2011). Nevertheless, the presence of residual TE insertions and low RIP signatures indicates that genome defence systems remain functional and may still influence local genomic regions. RIP-associated mutagenesis has been shown to contribute to gene diversification, pseudogenisation, and lineage divergence in fungi (van Wyk et al., 2019; Li et al., 2017). The low level of repetitive elements further supports the hypothesis that *F. avenaceum* is sexually active in nature, as active sexual cycles are typically associated with effective genome defence mechanisms such as RIP. Consistent with this, previous studies have reported that only ∼1% of the *F. avenaceum* genome consists of repetitive elements (Lysøe et al, 2014), a characteristic feature of fungi with active sexual reproduction and efficient suppression of TE accumulation.

Both Indian isolates showed relatively low TE content and RIP-associated genomic coverage compared with their respective reference groups. These comparative patterns may reflect differences in repeat abundance, sequence composition, or genomic context. However, RIP-associated coverage does not directly measure the current activity or efficiency of the RIP mechanism, and TE abundance alone cannot establish the historical rate of transposition or genome stability. Further analyses would be required to investigate the mechanisms underlying these patterns.

These genomic resources provide a foundation for investigating potential pathogenicity-associated genes and pathways. However, our ongoing transcriptome and dual RNA-seq studies involving diverse wheat genetic backgrounds are expected to clarify host–pathogen transcriptional responses and help prioritise candidates for subsequent functional validation.

This study presents hybrid genome assemblies for two Indian wheat-associated isolates, *F. graminearum* TNW1 and *F. avenaceum* DMW8, recovered from FHB-symptomatic fields in 2024-2025. The analyses identified shared predicted functional features, differences in secondary metabolite biosynthetic profiles, reduced CAZyme repertoires relative to their respective reference groups, low TE content, and contrasting gene family turnover. These findings provide genomic resources for further investigation of Fusarium associated with Indian wheat. Additional isolates, gene expression analyses, metabolite profiling, and functional assays will be needed to determine how the observed genomic differences relate to toxin production and wheat-pathogen interactions.

## Data availability

The genomic data, including raw Illumina NovaSeq 6000 and Oxford Nanopore PromethION sequence reads from two Indian isolates, *F. graminearum* TNW1 and *F. avenaceum* DMW8, have been deposited in the DDBJ/ENA/GenBank databases under BioProject accession number PRJNA1359619. Additional supporting data are available through https://data.mendeley.com/datasets/m2xy8zxx4g/1. All Shell, Python, and R scripts used for the analyses are available at https://github.com/govardhanchoppadandi/Fusarium-comparative-genomics, including the documented computational workflows and analysis pipelines used in this study.

## Conflict of Interest

The authors declare no conflicts of interest.

## Funding

Govardhan Choppadandi acknowledges the INSPIRE Fellowship (IF230586) awarded by the Department of Science and Technology (DST), Government of India, for financial support during his PhD research. Sudhir Navathe, Yashavanthakumar KJ and Govardhan Choppadandi acknowledge funding received from Agharkar research institute under project ARI-010.

## Authors contribution

**GC**: Execution: sequencing, isolation, manuscript writing, drafting, data analysis. **SN**: Conceptualisation, data analysis, project monitoring, fund acquisition, manuscript draft writing, and revisions. **Y. KJ**: Conceptualisation, project monitoring, manuscript revisions

## Acknowledgements

The authors gratefully acknowledge the MACS-Agharkar Research Institute, Pune, for providing the infrastructure.

---

**Figure S1:**
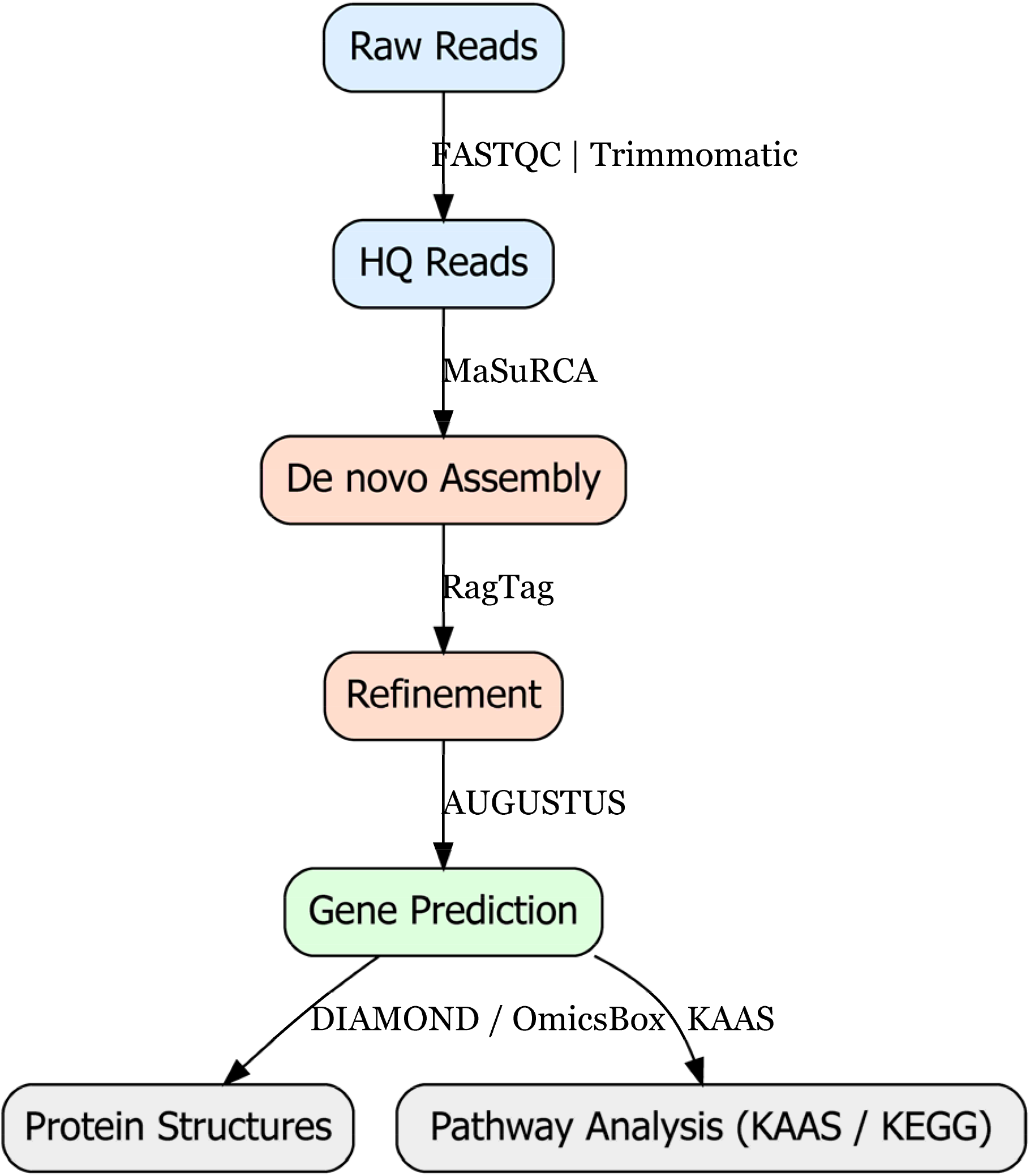
Genome assembly and annotation pipeline. Raw reads were processed, assembled, refined, and used for gene prediction, followed by functional and pathway annotation.

**Figure S2.**
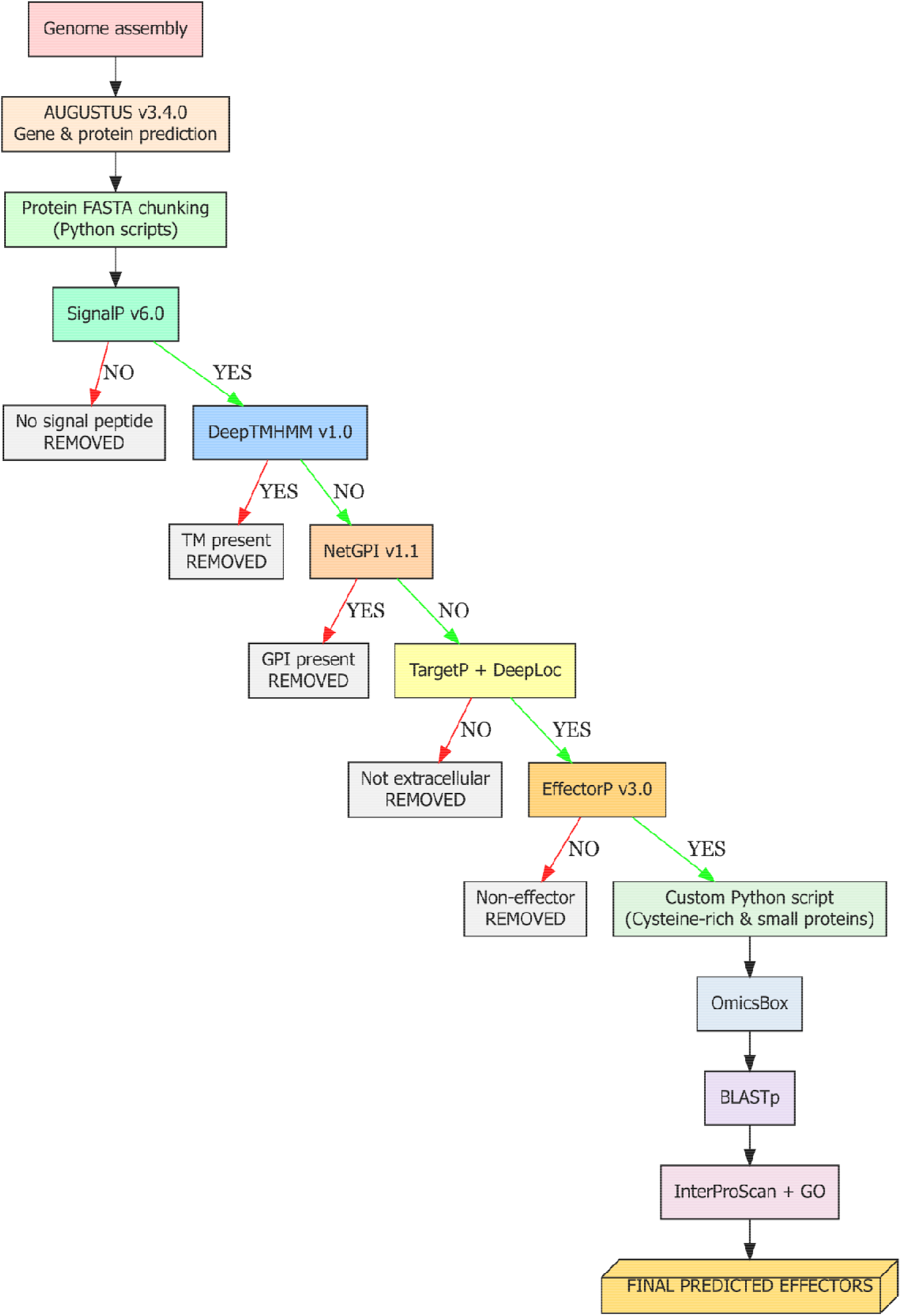
Effector prediction pipeline. Predicted proteins were filtered for secretion, transmembrane regions, GPI anchors, and extracellular localization, followed by effector prediction. Final candidates were refined by size and cysteine content and functionally annotated.

**Figure S3:**
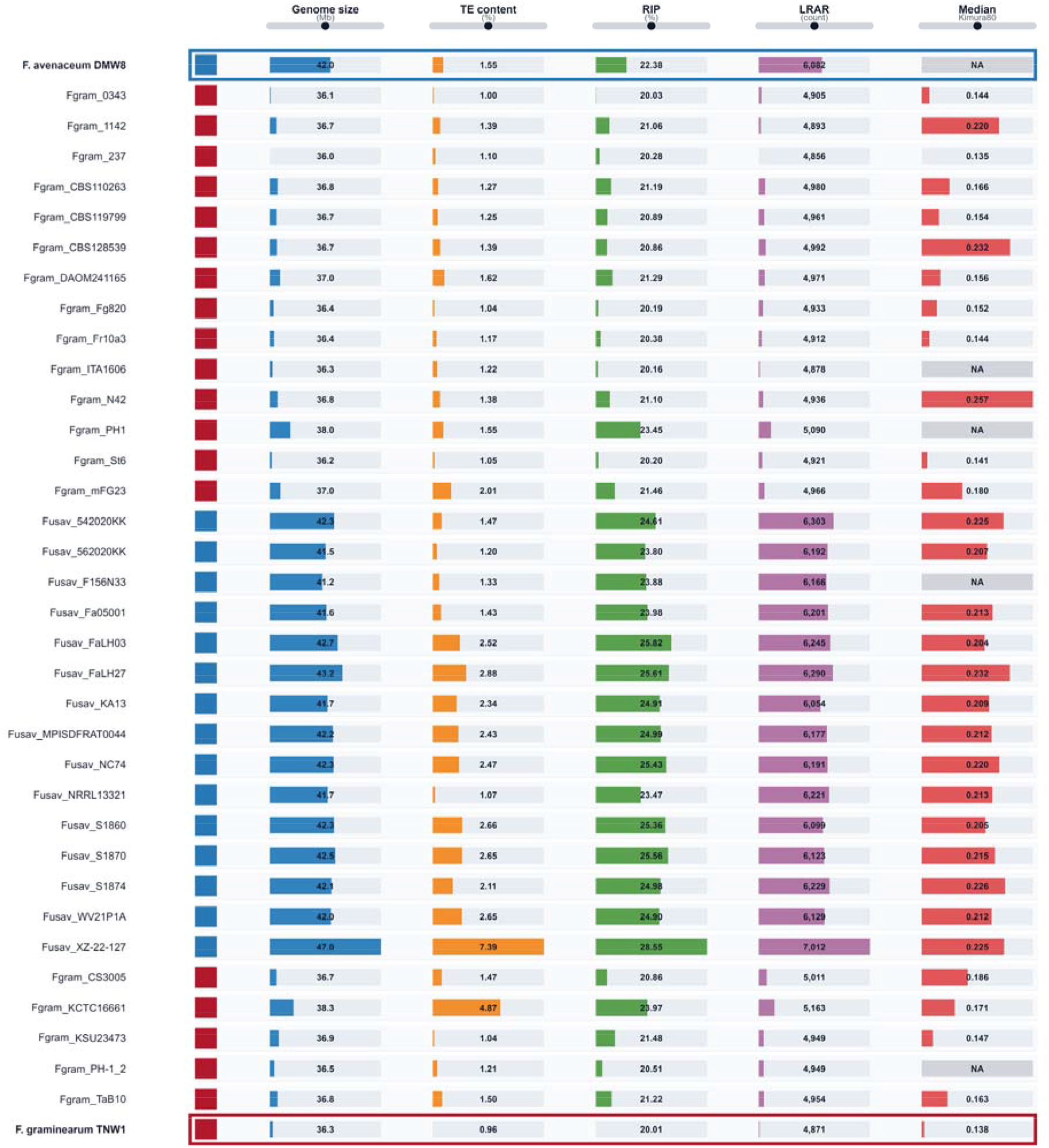
Integrated genomic landscape of 36 *Fusarium* genomes. Genome size, TE content, RIP, LRAR count, and median Kimura80 divergence are shown as aligned quantitative tracks. Blue and red indicate *F. avenaceum* and *F. graminearum*, respectively; DMW8 and TNW1 are highlighted, and missing Kimura80 values are shown as NA.

**Figure S4:**
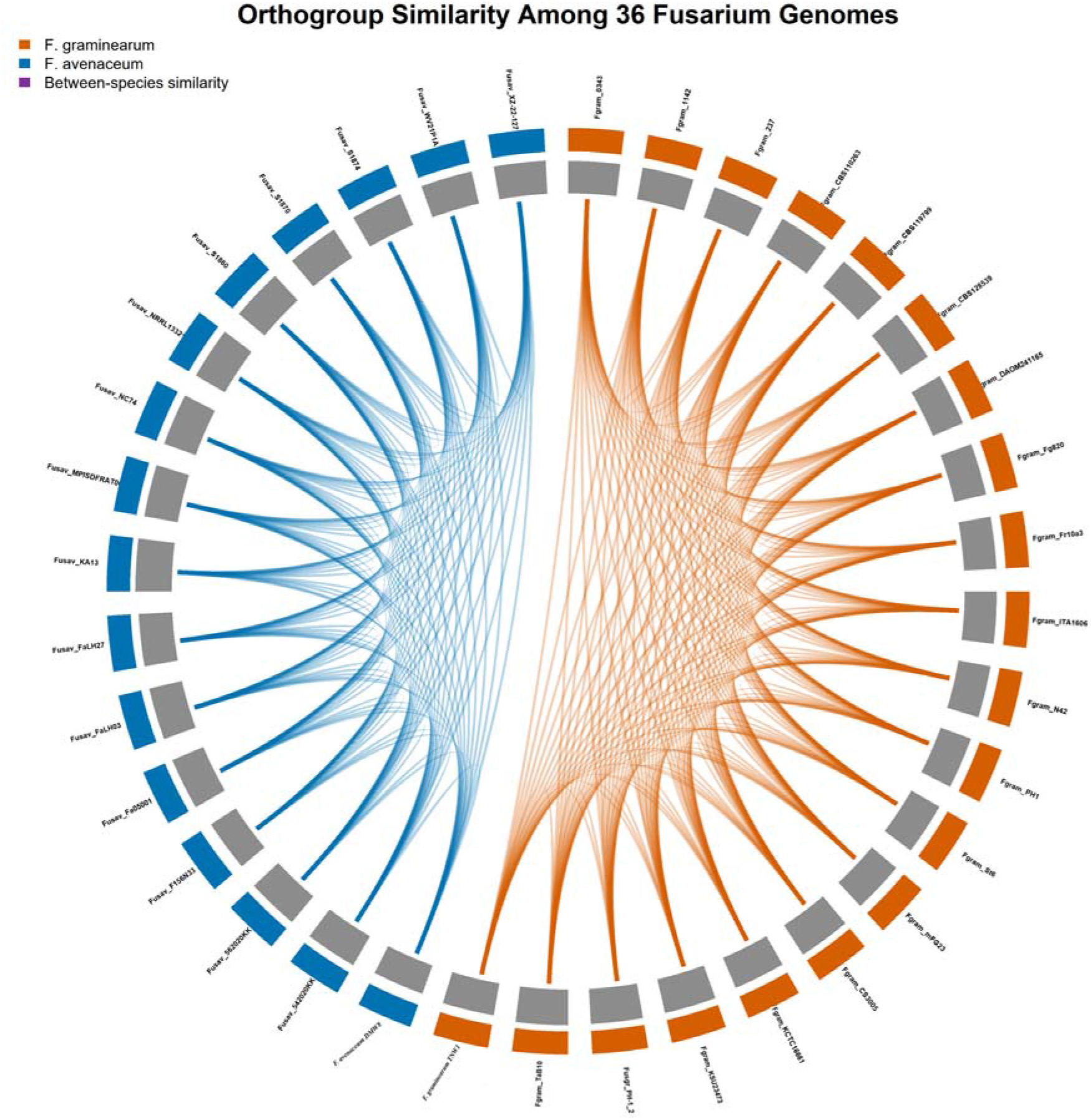
Orthogroup similarity among 36 *Fusarium* genomes. Orange and blue indicate *F. graminearum* and *F. avenaceum*, respectively. Connecting ribbons represent shared orthogroups, with colours indicating within-species similarity and between-species similarity.

**Figure S5:**
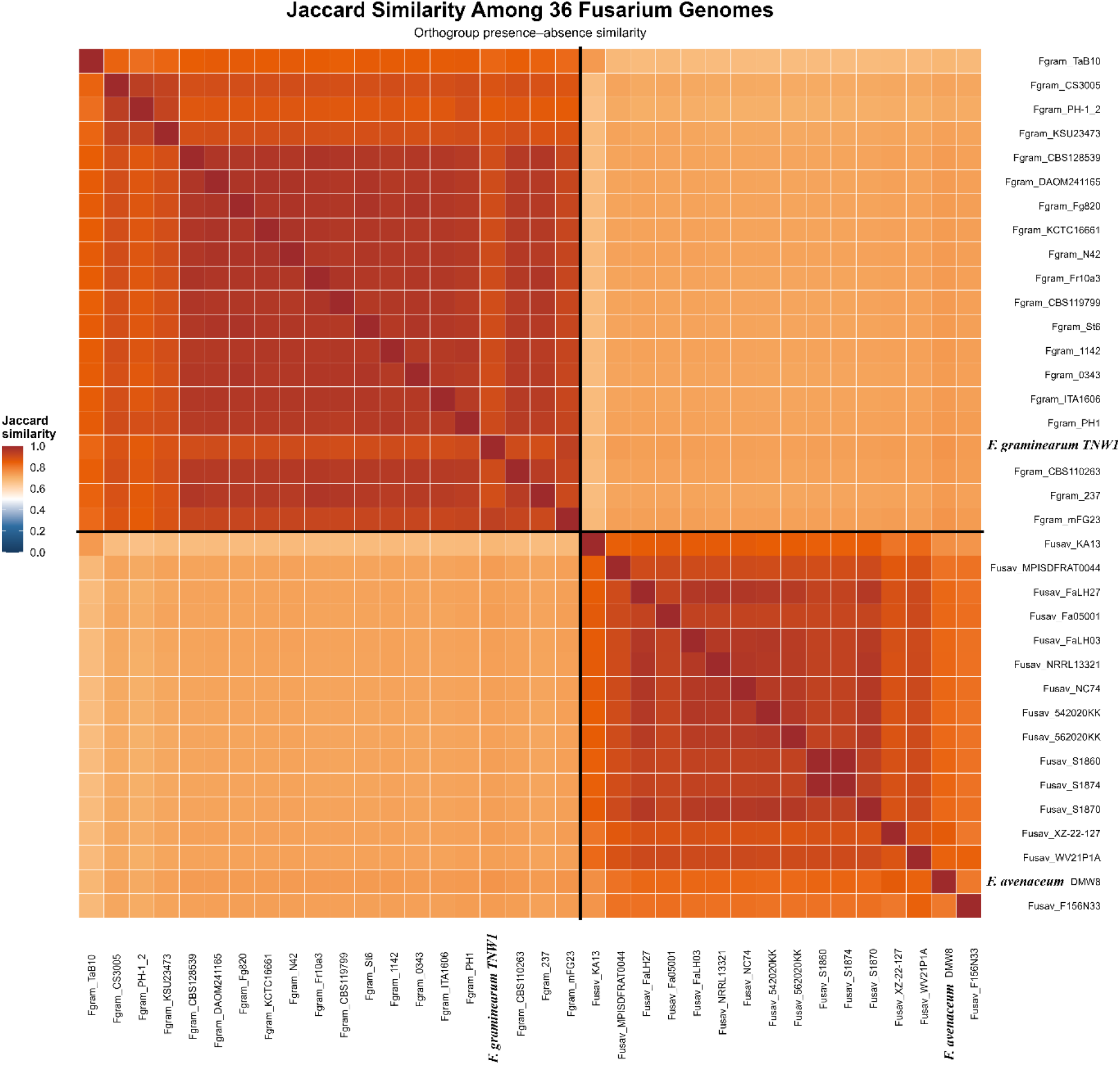
Pairwise Jaccard similarity among 36 *Fusarium* genomes. The heatmap shows orthogroup presence–absence similarity between genome pairs, with Jaccard similarity values ranging from 0 (no shared orthogroups) to 1 (identical orthogroup profiles). Black lines separate the *F. graminearum* and *F. avenaceum* genome groups.

## Notes

https://data.mendeley.com/datasets/m2xy8zxx4g/1.

https://github.com/govardhanchoppadandi/Fusarium-comparative-genomics

### Competing Interest Statement

The authors have declared no competing interest.

## References

Adrion, J.R, Song, M.J, Schrider, D.R, Hahn, M.W, Schaack, S, 2017. Genome-wide estimates of transposable element insertion and deletion rates in *Drosophila melanogaster*. Genome Biol. Evol. 9, 1329–1340. 10.1093/gbe/evx050.

Albalat, R., Cañestro, C, 2016. Evolution by gene loss. Nat. Rev. Genet. 17(7), 379–391. 10.1038/nrg.2016.39

Andrews, S, 2010. FastQC (Version 0.12.1) [software]. Babraham Bioinformatics, Cambridge, UK. Available at: http://www.bioinformatics.babraham.ac.uk/projects/fastqc/ (accessed 1 May 2026).

Armenteros, J.J.A, Salvatore, M, Emanuelsson, O, Winther, O, von Heijne, G, Elofsson, A, Nielsen, H, 2019. Detecting sequence signals in targeting peptides using deep learning. Life Sci. Alliance 2, e201900429. 10.26508/lsa.201900429.

Armer, V, Urban, M, Ashfield, T, Deeks, M.J, Hammond-Kosack, K.E, 2023. The trichothecene mycotoxin deoxynivalenol facilitates cell-to-cell invasion during wheat-tissue colonisation by Fusarium graminearum. bioRxiv [preprint]. 10.1101/2023.12.05.570169.

Badet, T, Croll, D, 2025. Phylogenomic signatures of repeat-induced point mutations across the fungal kingdom. PLoS Biol. 23, e3003433. 10.1371/journal.pbio.3003433.

Baril, T, Galbraith, J, Hayward, A, Grey, E, 2024. Earl Grey: a fully automated user-friendly transposable element annotation and analysis pipeline. Mol. Biol. Evol. 41, msae068. 10.1093/molbev/msae068.

Belyayev, A, 2014. Bursts of transposable elements as an evolutionary driving force. J. Evol. Biol. 27, 2573–2584. 10.1111/jeb.12513.

Blin, K, Shaw, S, Augustijn, H.E, Reitz, Z.L, Biermann, F, Alanjary, M, Fetter, A, Terlouw, B.R, Metcalf, W.W, Helfrich, E.J.N, van Wezel, G.P, Medema, M.H, Weber, T, 2023. antiSMASH 7.0: new and improved predictions for detection, regulation, chemical structures, and visualization. Nucleic Acids Res. 51, W46–W50. 10.1093/nar/gkad344.

Borowiec, M.L, 2016. AMAS: a fast tool for alignment manipulation and computing of summary statistics. PeerJ 4, e1660. 10.7717/peerj.1660.

Cantalapiedra, C.P, Hernández-Plaza, A, Letunic, I, Bork, P, Huerta-Cepas, J, 2021. eggNOG-mapper v2: functional annotation, orthology assignments, and domain prediction at the metagenomic scale. Mol. Biol. Evol. 38, 5825–5829. 10.1093/molbev/msab293.

Capella-Gutiérrez, S, Silla-Martínez, J.M, Gabaldón, T, 2009. trimAl: a tool for automated alignment trimming in large-scale phylogenetic analyses. Bioinformatics 25, 1972–1973. 10.1093/bioinformatics/btp348.

Castanera, R, López-Varas, L, Borgognone, A, LaButti, K, Lapidus, A, Schmutz, J, Grimwood, J, Pérez, G, Pisabarro, A.G, Grigoriev, I.V, Stajich, J.E, Ramírez, L, 2016. Transposable elements versus the fungal genome: Impact on whole-genome architecture and transcriptional profiles. PLoS Genet. 12(6), e1006108. 10.1371/journal.pgen.1006108.

Chan, P.P, Lin, B.Y, Mak, A.J, Lowe, T.M, 2021. tRNAscan-SE 2.0: improved detection and functional classification of transfer RNA genes. Nucleic Acids Res. 49, 9077–9096. 10.1093/nar/gkab688.

Chapman, B, Chang, J, 2000. Biopython: Python tools for computational biology. ACM SIGBIO Newsl. 20, 15–19. 10.1145/360262.360268.

Chen, J, Basting, P.J, Han, S, Garfinkel, D.J, Bergman, C.M, 2023. Reproducible evaluation of transposable element detectors with McClintock 2 guides accurate inference of Ty insertion patterns in yeast. Mob. DNA 14, 8. 10.1186/s13100-023-00296-4.

Chen, J, Wrightsman, T.R, Wessler, S.R, Stajich, J.E, 2017. RelocaTE2: a high resolution transposable element insertion site mapping tool for population resequencing. PeerJ 5, e2942. 10.7717/peerj.2942.

Chen, S, Zhou, Y, Chen, Y, Gu, J, 2018. fastp: an ultra-fast all-in-one FASTQ preprocessor. Bioinformatics 34, 884–890. 10.1093/bioinformatics/bty560.

Crous, P.W, Lombard, L, Sandoval-Denis, M, Seifert, K.A, Schroers, H.J, Chaverri, P, Gené, J, Guarro, J, Hirooka, Y, Bensch, K, Kema, G.H.J, Lamprecht, S.C, Cai, L, Rossman, A.Y, Stadler, M, Summerbell, R.C, Taylor, J.W, Ploch, S, Visagie, C.M, Thines, M, 2021. Fusarium: more than a node or a foot-shaped basal cell. Stud. Mycol. 98, 100116. 10.1016/j.simyco.2021.100116.

DeGenring, L.M, Gemechu, A, Rafiei, V, Broz, K, Regasa, G.H, Munsamy, K, Yilmaz, N, Dong, Y, Rouse, M.N, Drott, M.T, 2026. The 2022 *Fusarium* head blight outbreak in Ethiopia: Emerging pathogens, mixed mycotoxins, and interspecies interactions. Plant Dis. 110(3), 890–902. 10.1094/PDIS-01-25-0126-RE.

Dickson, J.G, 1942. Scab of wheat and barley and its control. US Department of Agriculture, Washington, DC, USA.

Dong, S, Raffaele, S, Kamoun, S, 2015. The two-speed genomes of filamentous pathogens: Waltz with plants. Curr. Opin. Genet. Dev. 35, 57–65. 10.1101/021774.

El-Sayed, R.A, Jebur, A.B, Kang, W, El-Demerdash, F.M, 2022. An overview on the major mycotoxins in food products: characteristics, toxicity, and analysis. J. Future Foods 2, 91–102. 10.1016/j.jfutfo.2022.03.002.

Emms, D.M, Kelly, S, 2019. OrthoFinder: phylogenetic orthology inference for comparative genomics. Genome Biol. 20, 238. 10.1186/s13059-019-1832-y.

Eskola, M, Kos, G, Elliott, C.T, Hajšlová, J, Mayar, S, Krska, R, 2019. Worldwide contamination of food-crops with mycotoxins: validity of the widely cited ‘FAO estimate’ of 25%. Crit. Rev. Food Sci. Nutr. 60, 2773–2789. 10.1080/10408398.2019.1658570.

Ferrigo, D, Raiola, A, Causin, R, 2016. Fusarium toxins in cereals: occurrence, legislation, factors promoting the appearance and their management. Molecules 21, 627. 10.3390/molecules21050627.

Fulnečková, J, Ševčíková, T, Fajkus, J, Lukešová, A, Lukeš, M, Vlček, Č, Lang, B.F, Kim, E, Eliáš, M, Sýkorová, E, 2013. A broad phylogenetic survey unveils the diversity and evolution of telomeres in eukaryotes. Genome Biol. Evol. 5, 468–483. 10.1093/gbe/evt019.

Gautier, C, Pinson-Gadais, L, Richard-Forget, F, 2020. Fusarium mycotoxins enniatins: an updated review of their occurrence, the producing Fusarium species, and the abiotic determinants of their accumulation in crop harvests. J. Agric. Food Chem. 68, 4788–4798. 10.1021/acs.jafc.0c00411.

Gíslason, M.H, Nielsen, H, Armenteros, J.J.A, Johansen, A.R, 2021. Prediction of GPI-anchored proteins with pointer neural networks. Curr. Res. Biotechnol. 3, 6–13. 10.1016/j.crbiot.2021.01.001.

Gladyshev, E, 2017. Repeat-induced point mutation and other genome defense mechanisms in fungi. Microbiol. Spectr. 5(4). 10.1128/microbiolspec.funk-0042-2017.

Hallgren, J, Tsirigos, K.D, Pedersen, M.D, Almagro Armenteros, J.J, Marcatili, P, Nielsen, H, Krogh, A, Winther, O, 2022. DeepTMHMM predicts alpha and beta transmembrane proteins using deep neural networks. bioRxiv [preprint]. 10.1101/2022.04.08.487609.

Han, S, Basting, P.J, Dias, G.B, Luhur, A, Zelhof, A.C, Bergman, C.M, 2021. Transposable element profiles reveal cell line identity and loss of heterozygosity in *Drosophila* cell culture. Genetics 219, iyab113. 10.1093/genetics/iyab113.

Hao, Z, Li, Y, Jiang, Y, Xu, J, Li, J, Luo, L, 2021. Genome sequence analysis of the fungal pathogen Fusarium graminearum using Oxford Nanopore technology. J. Fungi 7, 699. 10.3390/jof7090699.

Hegde, G.M, Devaiah, M, Uday, G, Patil, P.V, 2023. First report of *Fusarium poae* causing head blight of wheat in India. Plant Dis. 107, 3302. 10.1094/pdis-05-23-0879-pdn.

Hess, J, Skrede, I, Wolfe, B.E, LaButti, K, Ohm, R.A, Grigoriev, I.V, Pringle, A, 2014. Transposable element dynamics among asymbiotic and ectomycorrhizal *Amanita* fungi. Genome Biol. Evol. 6, 1564–1578. 10.1093/gbe/evu121.

Hubley, R, Finn, R.D, Clements, J, Eddy, S.R, Jones, T.A, Bao, W, Smit, A.F.A, Wheeler, T.J, 2016. The Dfam database of repetitive DNA families. Nucleic Acids Res. 44, D81–D89. 10.1093/nar/gkv1272.

Inglis, P.W, Pappas, M.D, Resende, L.V, Grattapaglia, D, 2018. Fast and inexpensive protocols for consistent extraction of high-quality DNA and RNA from challenging plant and fungal samples for high-throughput SNP genotyping and sequencing applications. PLoS One 13, e0206085. 10.1371/journal.pone.0206085.

Jestoi, M, 2008. Emerging Fusarium-mycotoxins fusaproliferin, beauvericin, enniatins, and moniliformin—a review. Crit. Rev. Food Sci. Nutr. 48, 21–49. 10.1080/10408390601062021.

Jones, P, Binns, D, Chang, H.-Y, Fraser, M, Li, W, McAnulla, C, McWilliam, H, Maslen, J, Mitchell, A, Nuka, G, Pesseat, S, Quinn, A.F, Sangrador-Vegas, A, Scheremetjew, M, Yong, S.-Y, Lopez, R, Hunter, S, 2014. InterProScan 5: genome-scale protein function classification. Bioinformatics 30, 1236–1240. 10.1093/bioinformatics/btu031.

Jørgensen, S.H, Frandsen, R.J.N, Nielsen, K.F, Lysøe, E, Søndergaard, T.E, Wimmer, R, Giese, H, Sørensen, J.L, 2014. *Fusarium graminearum* PKS14 is involved in orsellinic acid and orcinol synthesis. Fungal Genet. Biol. 70, 24–31. 10.1016/j.fgb.2014.06.008.

Käll, L, Krogh, A, Sonnhammer, E.L.L, 2007. Advantages of combined transmembrane topology and signal peptide prediction: the Phobius web server. Nucleic Acids Res. 35, W429–W432. 10.1093/nar/gkm256.

Katoh, K., Standley, D. M., 2013. MAFFT multiple sequence alignment software version 7: improvements in performance and usability. Mol. Biol. Evol. 30, 772–780. 10.1093/molbev/mst010.

Kaul, N, Kashyap, P.L, Kumar, S, Singh, D, Singh, G.P, 2022. Genetic diversity and population structure of head blight disease causing fungus *Fusarium graminearum* in northern wheat belt of India. J. Fungi 8, 820. 10.3390/jof8080820.

Kazan, K., Gardiner, D.M., Manners, J.M., 2012. On the trail of a cereal killer: recent advances in *Fusarium graminearum* pathogenomics and host resistance. Mol. Plant Pathol. 13, 399–413. 10.1111/j.1364-3703.2011.00762.x.

Keane, T.M., Wong, K., Adams, D.J., 2013. RetroSeq: transposable element discovery from next-generation sequencing data. Bioinformatics 29, 389–390. 10.1093/bioinformatics/bts697.

Kelkar, Y.D., Ochman, H., 2012. Causes and consequences of genome expansion in fungi. Genome Biol. Evol. 4(1), 13–23. 10.1093/gbe/evr124.

Kofler, R, Gómez-Sánchez, D, Schlötterer, C, 2016. PoPoolationTE2: comparative population genomics of transposable elements using Pool-Seq. Mol. Biol. Evol. 33, 2759–2764. 10.1093/molbev/msw137.

Kokkonen, M., Ojala, L., Parikka, P., Jestoi, M., 2010. Mycotoxin production of selected *Fusarium* species at different culture conditions. Int. J. Food Microbiol. 143, 17–25. 10.1016/j.ijfoodmicro.2010.07.015.

Kozlov, A.M., Darriba, D., Flouri, T., Morel, B., Stamatakis, A., 2019. RAxML-NG: a fast, scalable, and user-friendly tool for maximum likelihood phylogenetic inference. Bioinformatics 35, 4453–4455. 10.1093/bioinformatics/btz305.

Kseemann, T., 2014. Barrnap 0.9: Rapid ribosomal RNA prediction [software]. https://github.com/tseemann/barrnap.

Kubicek, C.P., Starr, T.L., Glass, N.L., 2014. Plant cell wall–degrading enzymes and their secretion in plant-pathogenic fungi. Annu. Rev. Phytopathol. 52, 427–451. 10.1146/annurev-phyto-102313-045831.

Li, H, Durbin, R, 2009. Fast and accurate short read alignment with Burrows–Wheeler transform. Bioinformatics 25, 1754–1760. 10.1093/bioinformatics/btp324.

Li, K, Yu, D, Yan, Z, Liu, N, Fan, Y, Wang, C, Wu, A, 2022. Exploration of mycotoxin accumulation and transcriptomes of different wheat cultivars during *Fusarium graminearum* infection. Toxins 14, 482. 10.3390/toxins14070482.

Li, L, Stoeckert, C.J, Roos, D.S, 2003. OrthoMCL: identification of ortholog groups for eukaryotic genomes. Genome Res. 13, 2178–2189. 10.1101/gr.1224503.

Li, W.C., Huang, C.H., Chen, C.L., Chuang, Y.C., Tung, S.Y., Wang, T.F., 2017. *Trichoderma reesei* complete genome sequence, repeat-induced point mutation, and partitioning of CAZyme gene clusters. Biotechnol. Biofuels 10, 1–20. 10.1186/s13068-017-0825-x.

Lo Presti, L., Lanver, D., Schweizer, G., Tanaka, S., Liang, L., Tollot, M., Zuccaro, A., Reissmann, S., Kahmann, R., 2015. Fungal effectors and plant susceptibility. Annu. Rev. Plant Biol. 66, 513–545. 10.1146/annurev-arplant-043014-114623.

Logrieco, A., Bottalico, A., Mulé, G., Moretti, A., Perrone, G., 2003. Epidemiology of toxigenic fungi and their associated mycotoxins for some Mediterranean crops. Eur. J. Plant Pathol. 109, 645–667. 10.1023/A:1026033021542.

Logrieco, A., Rizzo, A., Ferracane, R., Ritieni, A., 2002. Occurrence of beauvericin and enniatins in wheat affected by *Fusarium avenaceum* head blight. Appl. Environ. Microbiol. 68(1), 82–85. 10.1128/AEM.68.1.82-85.2002.

López Díaz, C., Ayhan, D.H., Rodríguez López, A., Gómez Gil, L., Ma, L.-J., Di Pietro, A., 2025. Transposons and accessory genes drive adaptation in a clonally evolving fungal pathogen. Nat. Commun. 16, 6982. 10.1038/s41467-025-62213-y.

Lysøe, E, Harris, L.J, Walkowiak, S, Subramaniam, R, Divon, H.H, et al, 2014. The genome of the generalist plant pathogen *Fusarium avenaceum* is enriched with genes involved in redox, signaling and secondary metabolism. PLoS One 9(11), e112703. 10.1371/journal.pone.0112703.

Mendes, F.K, Vanderpool, D, Fulton, B, Hahn, M.W, 2021. CAFE 5 models variation in evolutionary rates among gene families. Bioinformatics 36, 5516–5518. 10.1093/bioinformatics/btaa1022.

Minh, B.Q, Schmidt, H.A, Chernomor, O, Schrempf, D, Woodhams, M.D, von Haeseler, A, Lanfear, R, 2020. IQ-TREE 2: new models and efficient methods for phylogenetic inference in the genomic era. Mol. Biol. Evol. 37, 1530–1534. 10.1093/molbev/msaa015.

Moriya, Y., Itoh, M., Okuda, S., Yoshizawa, A., and Kanehisa, M. 2007. KAAS: an automatic genome annotation and pathway reconstruction server. Nucleic Acids Res. 35, W182–W185

Muszewska, A, Steczkiewicz, K, Stepniewska-Dziubinska, M, Ginalski, K, 2019. Transposable elements contribute to fungal genes and impact fungal lifestyle. Sci. Rep. 9(1). 10.1038/s41598-019-40965-0.

Nelson, M.G, Linheiro, R.S, Bergman, C.M, 2017. McClintock: an integrated pipeline for detecting transposable element insertions in whole-genome shotgun sequencing data. G3 (Bethesda) 7, 2763–2778. 10.1534/g3.117.043893.

Ødum, M.T, Teufel, F, Thumuluri, V, Almagro Armenteros, J.J, Johansen, A.R, Winther, O, Nielsen, H, 2024. DeepLoc 2.1: multi-label membrane protein type prediction using protein language models. Nucleic Acids Res. 52, W215–W220. 10.1093/nar/gkae237.

Platzer, A, Nizhynska, V, Long, Q, 2012. TE-Locate: a tool to locate and group transposable element occurrences using paired-end next-generation sequencing data. Biology 1, 395–410. 10.3390/biology1020395.

Pleadin, J, Frece, J, Markov, K, 2019. Mycotoxins in food and feed. Adv. Food Nutr. Res. 89, 297–345. 10.1016/bs.afnr.2019.02.007.

Stanke, M., Steinkamp, R., Waack, S., Morgenstern, B.: AUGUSTUS: a web server for gene finding in eukaryotes. Nucleic Acids Res. 32(Web Server issue), W309–W312 (2004). 10.1093/nar/gkh379

Quinlan, A.R, 2014. BEDTools: the Swiss-Army tool for genome feature analysis. Curr. Protoc. Bioinformatics 47, 11.12.1–11.12.34. 10.1002/0471250953.bi1112s47.

Raffaele, S, Kamoun, S, 2012. Genome evolution in filamentous plant pathogens: Why bigger can be better. Nat. Rev. Microbiol. 10(6), 417–430. 10.1038/nrmicro2790.

Rauwane, M.E, Ogugua, U.V, Kalu, C.M, Ledwaba, L.K, Woldesemayat, A.A, Ntushelo, K, 2020. Pathogenicity and virulence factors of *Fusarium graminearum* including factors discovered using next generation sequencing technologies and proteomics. Microorganisms 8, 305. 10.3390/microorganisms8020305.

Rhoades, N.A, McCormick, S.P, Vaughan, M.M, Hao, G, 2024. The emerging *Fusarium graminearum* NA3 population produces high levels of mycotoxins in wheat and barley. Toxins 16, 408. 10.3390/toxins16090408.

Robb, S.M.C, Lu, L, Valencia, E, Burnette, J.M, Okumoto, Y, Wessler, S.R, Stajich, J.E, 2013. The use of RelocaTE and unassembled short reads to produce high-resolution snapshots of transposable element generated diversity in rice. G3 (Bethesda) 3, 949–957. 10.1534/g3.112.005348.

Rouxel, T, Grandaubert, J, Hane, J.K, Hoede, C, van de Wouw, A.P, Couloux, A, Dominguez, V, Anthouard, V, Bally, P, Bourras, S, et al, 2011. Effector diversification within compartments of *Leptosphaeria maculans* genome affected by repeat-induced point mutations. Nat. Commun. 2, 1–10. 10.1038/ncomms1189.

Selvarajan, H, Los, L.D, Bakker, M.G, 2026. Fusarium graminearum. Trends Microbiol. 10.1016/j.tim.2026.01.007.

Singh, R.P, Hodson, D.P, Singh, P.K, Lan, C, He, X, Lagudah, E.S, Juliana, P, Ayliffe, M, Bhavani, S, Saunders, D.G.O, Huerta-Espino, J, 2025. Challenges to wheat disease resistance and current global strategies. Annu. Rev. Phytopathol. 63, 201–224. 10.1146/annurev-phyto-121923-082727.

Sperschneider, J, Dodds, P.N, 2022. EffectorP 3.0: prediction of apoplastic and cytoplasmic effectors in fungi and oomycetes. Mol. Plant Microbe Interact. 35, 146–156. 10.1094/MPMI-08-21-0201-R.

Stukenbrock, E.H, 2013. Evolution, selection and isolation: A genomic view of speciation in fungal plant pathogens. New Phytol. 199(4), 895–907. 10.1111/nph.12374.

Summerell, B.A, Salleh, B, Leslie, J.F, 2003. A utilitarian approach to Fusarium identification. Plant Dis. 87, 117–128. 10.1094/PDIS.2003.87.2.117.

Sun, J, Lu, F, Luo, Y, Bie, L, Xu, L, Wang, Y, 2023. OrthoVenn3: an integrated platform for exploring and visualizing orthologous data across genomes. Nucleic Acids Res. 51, W397–W403. 10.1093/nar/gkad313.

Sun, X, Yang, R, Tang, H, Ma, M, Chen, H, Chang, X, Zhang, M, Gong, G, 2025. Diversity and pathogenicity of Fusarium species associated with fusarium head blight in wheat and maize cropping systems in Sichuan Province. Sci. Rep. 15, 5984. 10.1038/s41598-024-83402-7.

Teufel, F, Almagro Armenteros, J.J, Johansen, A.R, Gíslason, M.H, Pihl, S.I, Tsirigos, K.D, Winther, O, Brunak, S, von Heijne, G, Nielsen, H, 2022. SignalP 6.0 predicts all five types of signal peptides using protein language models. Nat. Biotechnol. 40, 1023–1025. 10.1038/s41587-021-01156-3.

van Wyk, S, Harrison, C.H, Wingfield, B.D, de Vos, L, van der Merwe, N.A, Steenkamp, E.T, 2019. The RIPper: a web-based tool for genome-wide quantification of repeat-induced point (RIP) mutations. PeerJ 7, e7447. 10.7717/peerj.7447.

Wheeler, T.J, Eddy, S.R, 2013. nhmmer: DNA homology search with profile HMMs. Bioinformatics 29, 2487–2489. 10.1093/bioinformatics/btt403.

Wicker, T, Sabot, F, Hua-Van, A, Bennetzen, J.L, Capy, P, Chalhoub, B, Flavell, A, Leroy, P, Morgante, M, Panaud, O, Paux, E, SanMiguel, P, Schulman, A.H, 2007. A unified classification system for eukaryotic transposable elements. Nature Reviews Genetics 8 (12), 973–982. 10.1038/nrg2165.

Wise, K, Woloshuk, C, Freije, A, 2015. Fusarium head blight (head scab). Purdue Extension Publication BP-33-W. Purdue University, West Lafayette, IN, USA. https://www.extension.purdue.edu/extmedia/BP/BP-33-W.pdf (accessed 1 May 2026).

Xu, M, Wang, Q, Wang, G, Zhang, X, Liu, H, Jiang, C, 2022. Combatting fusarium head blight: advances in molecular interactions between Fusarium graminearum and wheat. Phytopathol. Res. 4, 37. 10.1186/s42483-022-00142-0.

Xu, Z, Wang, H, 2007. LTR_FINDER: an efficient tool for the prediction of full-length LTR retrotransposons. Nucleic Acids Res. 35, W265–W268. 10.1093/nar/gkm286.

Yu, T, Huang, X, Dou, S, Tang, X, Luo, S, Theurkauf, W.E, Lu, J, Weng, Z, 2021. A benchmark and an algorithm for detecting germline transposon insertions and measuring de novo transposon insertion frequencies. Nucleic Acids Res. 49, e44. 10.1093/nar/gkab010.

Zheng, J, Ge, Q, Yan, Y, Zhang, X, Huang, L, Yin, Y, 2023. dbCAN3: automated carbohydrate-active enzyme and substrate annotation. Nucleic Acids Res. 51, W115–W121. 10.1093/nar/gkad328.

Zhuang, J, Wang, J, Theurkauf, W, Weng, Z, 2014. TEMP: a computational method for analyzing transposable element polymorphism in populations. Nucleic Acids Res. 42, 6826–6838. 10.1093/nar/gku323.

Tegenfeldt, F., Kuznetsov, D., Manni, M., Berkeley, M., Zdobnov, E.M., Kriventseva, E.V.: OrthoDB and BUSCO update: annotation of orthologs with wider sampling of genomes. Nucleic Acids Res. (2025). 10.1093/nar/gkae987

